# Intermittent KHz-frequency electrical stimulation selectively engages small unmyelinated vagal afferents

**DOI:** 10.1101/2021.01.30.428827

**Authors:** Yao-Chuan Chang, Umair Ahmed, Naveen Jayaprakash, Ibrahim Mughrabi, Qihang Lin, Yi-Chen Wu, Michael Gerber, Adam Abbas, Anna Daytz, Arielle H. Gabalski, Jason Ashville, Socrates Dokos, Loren Rieth, Timir Datta-Chaudhury, Kevin Tracey, Tianruo Guo, Yousef Al-Abed, Stavros Zanos

## Abstract

Afferent and efferent vagal fibers mediate bidirectional communication between the brain and visceral organs. Small, unmyelinated C-afferents constitute the majority of vagal fibers, play critical roles in numerous interoceptive circuits and autonomic reflexes in health and disease and may contribute to the efficacy and safety of vagus nerve stimulation (VNS). Selective engagement of C-afferents with electrical stimuli has not been feasible, due to the default fiber recruitment order: larger fibers first, smaller fibers last. Here, we determine and optimize an electrical stimulus that selectively engages vagal C-afferents. Intermittent KHz-frequency electrical stimulation (KES) activates motor and, preferentially, sensory vagal neurons in the brainstem. During KES, asynchronous activity of C-afferents increases, while that of larger fibers remains largely unchanged. In parallel, KES effectively blocks excitability of larger fibers while moderately suppressing excitability of C-afferents. By compiling selectivity indices in individual animals, we find that optimal KES parameters for C-afferents are >5KHz frequency and 7-10 times engagement threshold (×T) intensity in rats, 15-25×T in mice. These effects can be explained in computational models by how sodium channel responses to KES are shaped by axonal size and myelin. Our results indicate that selective engagement of vagal C-afferents is attainable by intermittent KES.

## Introduction

Homeostasis in organisms is maintained through orchestrated operation of immune, endocrine and neural processes. The autonomic nervous system (ANS) plays central role in homeostatic control, through an interconnected network of fast-acting reflexes that regulate the function of internal organs in real time. Autonomic reflexes comprise sensing and effector mechanisms in body organs, integrator systems in the brain, and peripheral nerves, which convey information between them. The vagus nerve is the main neural conduit of body-brain communication, mediating bidirectional transmission of sensory (afferent) and motor (efferent) information. The vast majority of nerve fibers in the vagus are small, unmyelinated, slowly conducting, C-type afferents(1). Vagal C-afferents mediate numerous and diverse functions, including sensing of nutrients(2) and regulation of appetite and glucose metabolism(3), effects of gut microbiome on brain function and cognition(4), neural regulation of breathing(5), modulation of immune responses to lung infections(6), and shaping of emotional responses by bodily feedback(7). Vagal C-afferents also constitute the afferent arm of cardiovascular reflexes(8), neuroimmune circuits in the gut(9), and the inflammatory reflex itself(10), a neuroimmune circuit that maintains immunological homeostasis throughout the body(11). Engagement of C-afferents by vagus nerve stimulation (VNS) may have therapeutic implications for neurostimulation-based therapies of arthritis(12), inflammatory bowel disease(13, 14), heart failure(15) and obesity(16).

Controlled, selective engagement of distinct nerve fiber types, separately from other fiber populations in the same nerve, is required for the study of their physiological and translational roles(17). Selective activation of C-afferents in the vagus is possible using optogenetic nerve stimulation(18) but that is currently only practical with mice in acute experiments, with limited value in preclinical models of chronic disease and unclear clinical applicability. On the other hand, electrical stimulation of the vagus can be delivered acutely or chronically, in various animal models, including mice(19), and in humans. However, there is no known electrical stimulus that selectively activates vagal C-fibers. That is mainly because of the natural recruitment order of nerve fibers: larger fibers (A- and B-type) are activated well before smaller fibers can be engaged(20), and at high stimulus intensities, required to activate C-fibers, larger fibers are also maximally activated(21, 22). The lack of a C-fiber-selective electrical stimulus hinders the study of the many interoceptive functions and autonomic reflexes in which C-afferents are involved, the translational testing of VNS in animal models of disease and, ultimately, the therapeutic potential of VNS.

Kilohertz-frequency electrical stimulation (KES) blocks nerve conduction (23-26) and is used in the vagus nerve for treating obesity(27) and in somatic sensory nerves for treating pain(28, 29), with an assumed mechanism of action that involves blocking of C-afferents. Using electrical and optogenetic stimulation, imaging, physiological and computational methods, we show here that KES of the vagus nerve can instead preferentially activate C-afferents while simultaneously blocking larger fibers, in a reliable and reversible manner. We also describe a method to select optimal KES frequency and intensity for C-afferent fiber activation in real time, in individual subjects.

## Results

### Intermittent KES activates motor and, preferentially, sensory vagal neurons in the brainstem

KES has been shown to block nerve fibers of different sizes(30), even though its mechanism of action and its effect on the fiber-associated neurons themselves remain elusive(26, 31). **To determine the effect of KES delivered to the cervical vagus on the activation level of neurons associated with different fiber types, we quantified a marker of neuronal activation**, c-Fos expression, in sensory and motor vagal neuronal populations in the brainstem. In anesthetized rats, we delivered sham or intermittent VNS (10 s on, 50 s off) for 30 minutes, comprising either KES trains (8-kHz frequency, 40µs pulse width, 2mA intensity) or comparable trains of “standard VNS” (30 Hz, 40µs, 2mA) (Figure 1A). We then counted c-Fos^+^ neurons in the nucleus tractus solitarius (NTS), a sensory region receiving projections primarily from C-afferents via the nodose ganglion(32, 33), and in the dorsal motor nucleus of the vagus (DMV), a motor region with cholinergic (ChAT^+^) cells providing efferent, mostly Aα- and B-fiber fibers, to the vagus(34) (Figure 1B). Sham VNS was associated with minimal c-Fos expression in NTS and DMV (Figure 1B-a). To our surprise, after 30 min of intermittent KES, we observed increased, compared to sham, c-Fos expression, stronger in the ipsilateral (to KES) sensory, NTS region, and weaker in the motor, DMV region (Figure 1B-b). As expected, 30-Hz VNS induced strong c-Fos expression in both ipsilateral (to VNS) sensory, NTS, and motor, DMV, regions (Figure 1B-c). Overall, KES induces a 1.7-fold increase, compared to sham, in c-Fos expression in NTS, with standard VNS producing a comparable 2.2-fold increase (Figure 1C); at the same time, KES induces a non-significant (0.9-fold) increase in c-Fos in DMV, much smaller than that induced by standard VNS (2.4-fold) (Figure 1C-D). This preferential activation of NTS over DMV by KES is demonstrated by a sensory neuronal-c-Fos^+^ selectivity index (NcSI), defined as c-Fos^+^ cell count ratio of NTS to DMV (Figure 1F) (Suppl. Figure S1D). Counts of c-Fos^+^ neurons in the sham stimulation condition are moderately greater than naïve animals in NTS, and no different in DMV (Figure 1C and 1D, and Suppl. Figure S1). Interestingly, 30Hz VNS causes a moderate increase of c-Fos^+^ cells in contralateral NTS and DMV (0.85- and 1.1-fold, respectively), whereas KES did not significantly affect contralateral c-Fos expression (Suppl. Figure S1B-D), suggesting that KES elicits a more “lateralized” neuronal activation.

**Figure 1:**
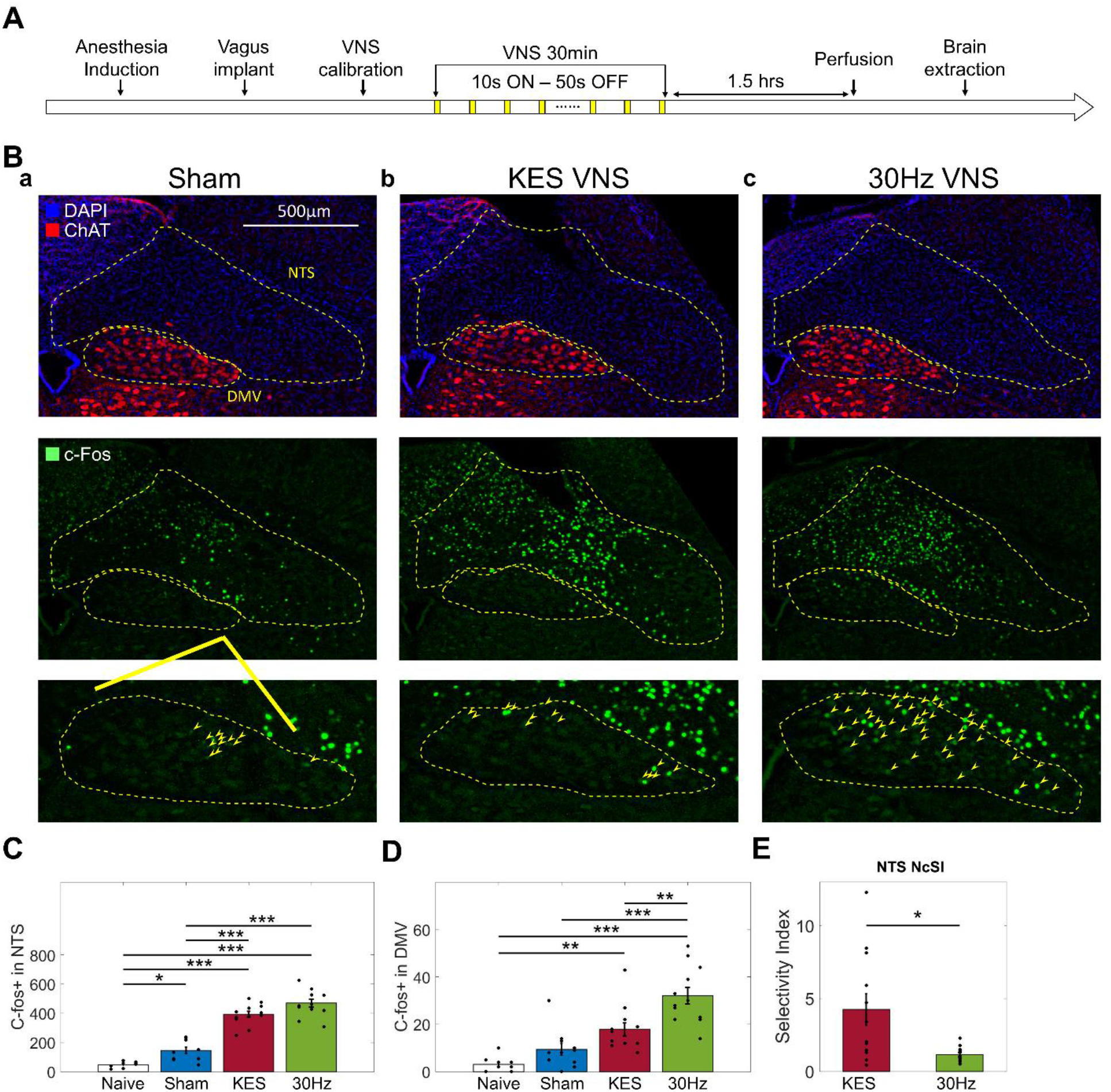
Intermittent kHz-frequency electrical stimulation (KES) activates motor and, preferentially, sensory vagal neurons in the brainstem. (A) Time course of experiments to quantify c-Fos-expressing neurons in the brainstem after VNS. (B) Representative immunohistochemistry images of sections across ipsilateral, to VNS, sensory and motor brainstem regions (yellow contours): nucleus tractus solitaries (NTS) and dorsal motor nucleus of the vagus (DMV), identified by DAPI (blue) and ChAT(red) (1^st^ row), each stained for c-Fos (green) (2^nd^ row). The insets (3^rd^ row) show ipsilateral DMV at higher magnification; arrows point to cells positive for c-Fos. (C) c-Fos^+^ cell numbers (mean ± SE) in ipsilateral NTS, in different groups of animals: naïve (white), sham stimulation (blue), KES (8-kHz, dark red), 30 Hz VNS (light green). Statistical comparisons between groups use one-way ANOVA and Tukey’s post-hoc tests (*p<0.05, **p<0.005, **p<0.0005). (D) Same as (C) but in ipsilateral DMV. (E) NTS neuronal-c-Fos^+^ selectivity index (NcSI) (mean ± SE), calculated as the fold change of c-Fos^+^ expression, with respect to sham average, in the NTS region over the one in DMV region, for the KES VNS group of animals and the 30 Hz group. Statistical comparison between groups uses 2-sample t-test (p<0.05).

### KES differentially affects asynchronous activity and excitability of vagal fiber types

**Our finding that KES increases activity of motor and, preferentially, sensory vagal neurons in the brainstem is in disagreement with the widely-held assumption that KES blocks nerve conduction(35, 36), even though that has recently been a matter of debate(31). To investigate whether fiber engagement by KES can explain this finding, we assessed changes in ongoing (asynchronous) activity of different fiber types during nerve stimulation**. Short (10 s-long) trains of KES (8-kHz, at a range of intensities) were delivered in the vagus of anesthetized rats while recording several physiological parameters and nerve potentials (Figure 2A). Changes in laryngeal EMG, HR and BI during VNS are real-time physiological responses driven by and correlating with asynchronous activity in A-, B- and C- fibers, respectively (Suppl. Figure S2), due to temporal summation of post-synaptic potentials(37). Likewise, optogenetic stimulation of cholinergic, B-fibers in ChAT-ChR mice causes bradycardia, whereas stimulation of glutamergic, C-fibers in VGluT-ChR mice slows down breathing, in a dose-dependent manner (Figure 2B). We used those responses to assess changes in asynchronous fiber activity elicited by VNS(37), since electrical artifacts during KES trains preclude direct recording of fiber potentials from the nerve. At low intensities, KES elicits A-fiber-associated EMG (Figure 2C; 2D-a; Suppl. Video 1D) and B-fiber associated HR responses; with increasing intensity, those responses are progressively suppressed and almost completely abolished at intensities >6-7 times threshold intensity (×T) (Figure 2C; 2D-a; Suppl. Video 1E). In contrast, C-fiber-associated BI responses appear at intensities >6-7×T, continue increasing up to 15×T and are eventually blocked above 30×T (Figure 2C; 2D-a; Suppl. Video 1F). These results suggest that within an intensity window of 7-15×T, asynchronous activity of C-fibers remains robust, whereas that of A- and B-fibers is blocked. These findings were replicated in mice, with a C-fiber intensity window of 10- 30×T (Suppl. Figure S3).

**Figure 2:**
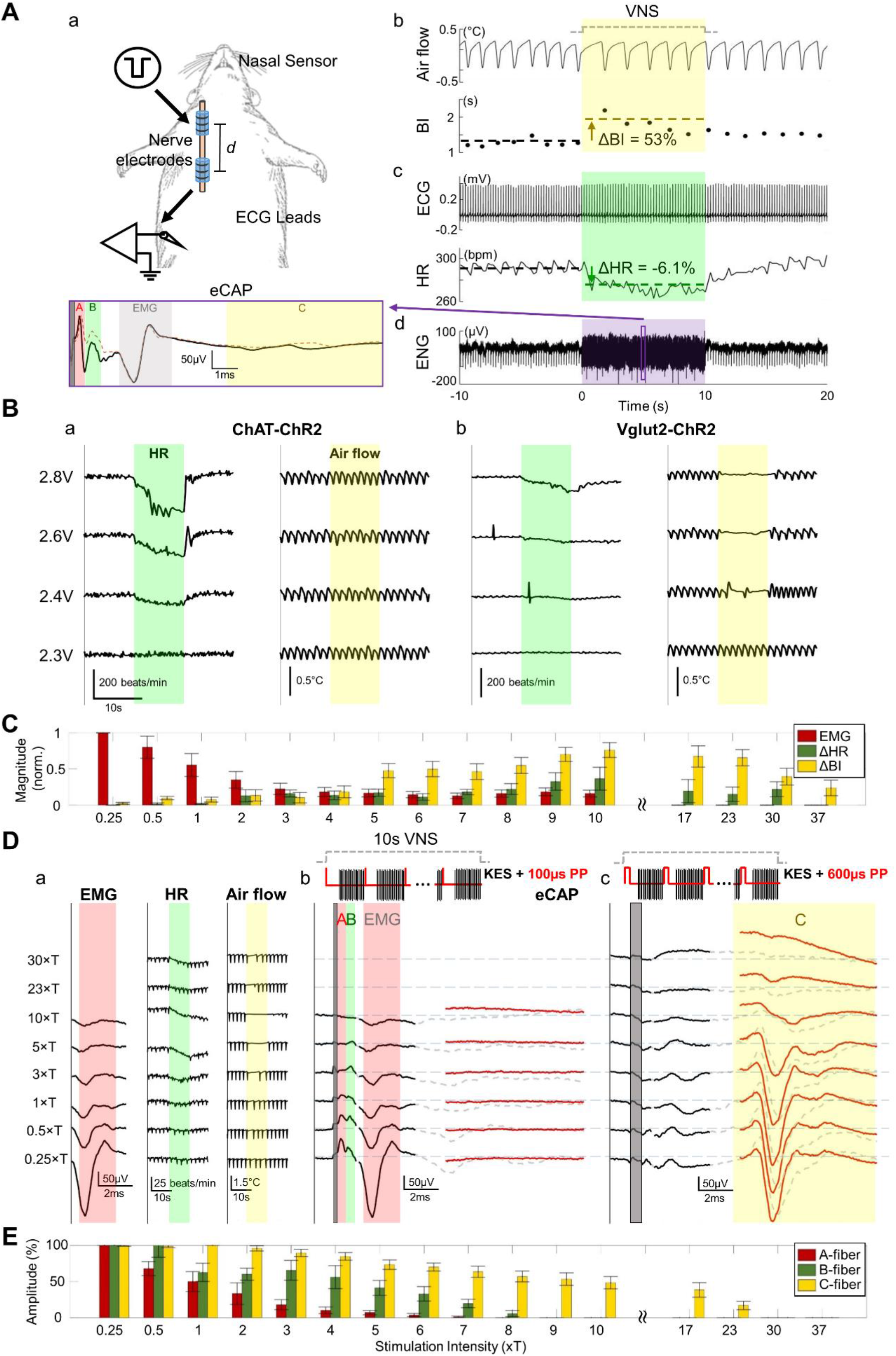
KES frequency and intensity interact to differentially control asynchronous activity and excitability of vagal fibers. (A) Schematic of a typical physiology experiment. Nerve electrodes are placed on the cervical vagus nerve for stimulation and recording of nerve activity; a nasal sensor and ECG leads record air flow and ECG, respectively (panel a). During electrical stimulation of the vagus (VNS), changes in breathing interval (BI), typically slowing of breathing or apnea (panel b), and in heart rate (HR), usually bradycardia (panel c), are observed. Single stimuli evoke compound action potentials (eCAPs), extracted from electroneurogram (ENG) (panel d), with early, intermediate and late components representing evoked volleys in A-, B- and C-fibers, respectively(37). (B) Optogenetic VNS delivered to VGluT-ChR transgenic mice causes slowing of breathing (panel a), whereas when delivered to ChAT-ChR mice causes bradycardia (panel b), in a dose-dependent manner. (C) Mean (±SE, N=4 animals) of normalized magnitude of physiological responses (EMG, red; ΔHR, green; ΔBI, yellow), elicited by KES of different intensities. (Linear Regression, *p<0*.*05* for intensity, across all physiological responses (EMG, HR, BI)). (D) Representative physiological responses elicited with KES (8-kHz, 970ms ON, 30ms OFF, 10s); as KES intensity increases (from bottom to top), (panel a) suppressed EMG, robust ΔBI with minimal ΔHR responses are observed; (panel b) eCAPs encompassing A- and B-fiber responses were compiled by delivering, intermittently with KES, single “probing” pulses (PP) of 100 us PW during the OFF window (every 1 s, for 10 s). A- or B-fiber evoked activity is mostly blocked at KES intensities above ∼5×T. C-fibers are not evoked with short probing pulses. (panel c) Same as (b), but this time eCAPs triggered with 600 µs-long probing pulses, to evoke C-fiber activity. In contrast to A- and B-, evoked C-fiber activity is maintained at KES intensities 5-10×T and progressively disappears at higher intensities. (E) Mean (±SE, N = 4 animals) of normalized amplitude of A-, B-, C-fiber evoked activity (red, green and yellow bars, respectively), for KES of different intensities; 100 µs probing pulses used for A- and B-fibers, 600 µs probing pulses used for C-fibers (ANOVA, *p<0*.*05* for intensity, across A-, B- and C-fiber responses).

**Our finding that KES increases markers of asynchronous activity of C-afferents does not agree with previous reports of C-afferent block by KES(35, 36)**.To investigate this apparent discrepancy, we assessed changes in fiber excitability during VNS by delivering single “probing” pulses, every 1 s, throughout the 10-s KES trains and measured fiber-specific, synchronous eCAPs(38) (Figure 2D-b and c). Addition of probing pulses does not alter the physiological responses to KES (Suppl. Figure S4). Compared to pre-stimulation levels, eCAP amplitude progressively decreases as KES intensity increases (Figure 2D-b and c). This suppression of excitability occurs almost immediately upon KES delivery (Suppl. Figure S5, 2^nd^ eCAP) and affects all fiber types. However, whereas A- and B- fiber excitability is almost completely abolished at intensities >7-8×T, C- fiber excitability gets suppressed at a much slower rate and C-fibers are still significantly excitable at intensities 7-20×T (Figure 2E). Importantly, A-, B- and C-fiber excitability returns to pre-stimulation levels after the end of each KES train (Suppl. Figure S5), suggesting that KES within this intensity range (<20×T for rats, <30×T for mice) does not produce irreversible effects.

### Selection of KES frequency and intensity optimize C-afferent selectivity

**Having demonstrated that KES selectively activates C-afferents in a frequency- and intensity-dependent manner, we next sought to experimentally determine the KES parameters that maximize such selectivity**. KES at relatively low frequencies (e.g. 1-kHz) elicits similar HR and BI responses to those elicited by 30-Hz trains with matched duration, intensity and PW (Suppl. Figure S13B), indicating limited selectivity for C-afferents. At higher frequencies (>5-kHz), high intensity KES results in similar BI responses as 30-Hz trains but with minimal HR responses, indicating selective activation of C-afferents (Figure 3A and Suppl. Figure S13A). Overall, the higher the frequency of KES, the smaller the HR effect is for a similar, to 30-Hz VNS, BI response (Figure 3B). In rats, selectivity for C-afferents is maximized at KES frequencies of 5-kHz or above, at intensities 8-10×T (Figure 3C). In experiments in mice, KES at intensities 15-25×T elicited similar BI responses as 30 Hz VNS, but with a much smaller HR response (Suppl. Figure S3).

**Figure 3:**
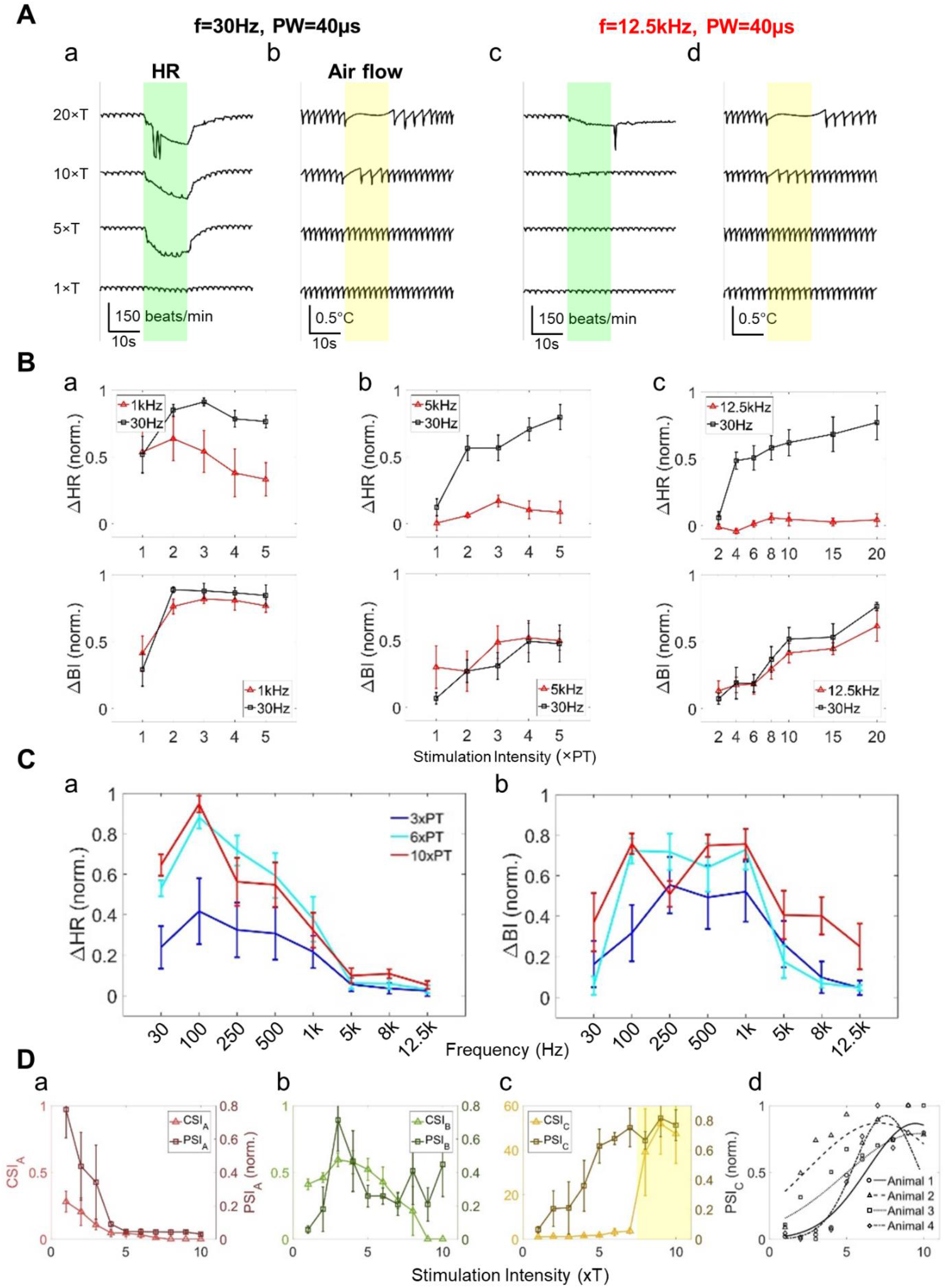
VNS frequencies over 5kHz at high intensities convey increased C-fiber selectivity. (A) Representative heart rate (HR) and breathing responses (airflow) elicited by stimuli of 12.5-kHz (40 µs PW, panel c and d), next to their 30 HZ, PW- and intensity-matched controls from the same animal (panel a and b). The HR responses are highly suppressed by 12.5-kHZ stimuli even with high VNS intensity, compared with 30Hz, whereas the BI responses remain similar. (B) Mean (±SE, N = 5 animals) normalized ΔHR and ΔBI responses for 1kHz (PW=500µs, panel a), 5kHz (PW=100µs, panel b), 12.5-kHz (PW=40µs, panel c) frequency stimuli (red curves) along with responses to their corresponding 30 Hz controls (black curves), as a function of stimulus intensity respectively. (ANCOVA, ΔHR: *p<0*.*05* for all pair-wise stimuli and intensity, and their interaction; ΔBI: *p>0*.*05* for all pair-wise stimuli and *p<0*.*05* intensity, *p<0*.*05* for stimuli/intensity interaction). (C) panel a: Mean (±SE, N = 5 animals) normalized ΔHR responses to trains of stimuli of varying frequencies (intensity ranging from 3×PT to 10×PT, shown in different color curves) and of identical PW (40 µs), as a function of frequency. (ANCOVA, *p<0*.*05* for frequency and intensity, and *p<0*.*05* for interaction). panel b: Same as (C) but for normalized ΔBI responses. (ANCOVA, *p<0*.*05* for frequency and intensity, and *p<0*.*05* for interaction). (D) Mean (±SE, N=4 animals) of CSI and normalized PSI values for A- (panel a), B- (panel b) and C-fibers (panel c) as function of KES (8-kHz) intensity (ANCOVA, *p<0*.*05* for all types of CSI and intensity, and their interaction; *p<0*.*05* for all types of PSI, and PSI/intensity interaction). KES of high intensity produces highest selectivity for C-fibers (yellow window in panel c). (panel d) C-fiber PSI values at different KES intensities and Gaussian fits in individual animals. Average RMSE for fits: 0.169.

Given the non-monotonic relationship between KES frequency, intensity and fiber selectivity, and the variability of this relationship between animals (Figure 3D), it would be desirable to determine optimal KES parameters for engaging a fiber type on an individual animal basis, in real-time. To personalize KES parameters for optimal fiber selectivity we measured fiber-specific responses to stimuli to compile physiological (PSI) or eCAP (CSI) fiber selectivity indices (Suppl. Figure S6C) (Equations given in Supplement). CSI and PSI provide subject-specific read-outs of fiber engagement to single trains of stimuli, as they both rely on acute physiological responses to stimulation(37, 39); PSI, in particular, is compiled from physiological markers that can be measured non-invasively in animals or humans(37). When applied to 30-Hz VNS trains with short-square pulses (100 μs), or other waveforms delivered at low intensity (1-3×T), PSI and CSI are maximal for A-fibers (Suppl. Figure S6-S10 for rats, and Suppl. Figure S11-12 for mice), which is expected given the low activation threshold of A-fibers(40). For 30-Hz VNS with long-square (>500 μs) or quasi-trapezoidal pulses at intermediate intensity, indices are maximal for B-fibers (Suppl. Figure S6-S10 for rats, and Suppl. Figure S11-S12 for mice), a finding that is also expected due to anodal block of A-fibers(40-42). When applied to KES trains, PSI and CSI for each of the 3 fiber types behave similarly (Figure 3D-, a-c). For C-fibers in particular, selectivity indices are maximal at relatively high KES intensities (Figure 3D-c). Even though KES intensity for optimal selectivity in engaging C-fibers is different between animals, it is always within the predicted intensity window (Figure 3D-d and Suppl. Figure S3D).

### Possible cell membrane mechanism of KES selectivity for C-afferents

**To mechanistically understand the effects of KES on fiber engagement at the level of the cell membrane, we simulated voltage responses to KES trains** in larger, myelinated fibers (A/B-type, 2.6 μm diameter) and in smaller, unmyelinated C-afferents (1.3 μm). At low KES frequencies (<1-kHz), both fiber types are activated with no selectivity (Figure 4A, a-c). At higher KES frequencies (>2-kHz), large fibers are blocked at low intensities, whereas C-afferents are progressively activated at increasingly higher intensities (Figure 4A, d-g) (Figure 4B), in agreement with the stimulus intensity window we observe experimentally. This fiber selectivity of high-frequency KES can be explained by the different activation and inactivation dynamics of sodium currents, which are present in both fiber types. At low intensities, action potentials (APs) are elicited in large fibers (Figure 4B-a1) as activation (m) and inactivation (h) gates are fully functional in a physiological range, while small fibers are unresponsive (Figure 4B-b1). At intermediate intensities, large fibers are quickly blocked, as the activation and inactivation gates start to passively follow the extracellular voltage changes (Figure 4B-a2), while small fibers remain unresponsive (Figure 4B-b2). At high intensities, APs are now elicited in small fibers as activation and inactivation gates become functional (Figure 4B-b3). Large, myelinated fibers are “tuned” to low KES intensities, large unmyelinated fibers to intermediate intensities and small, unmyelinated fibers to high intensities (Figure 4C), suggesting that both the presence of myelin and the size of fibers shapes the responses to KES.

**Figure 4:**
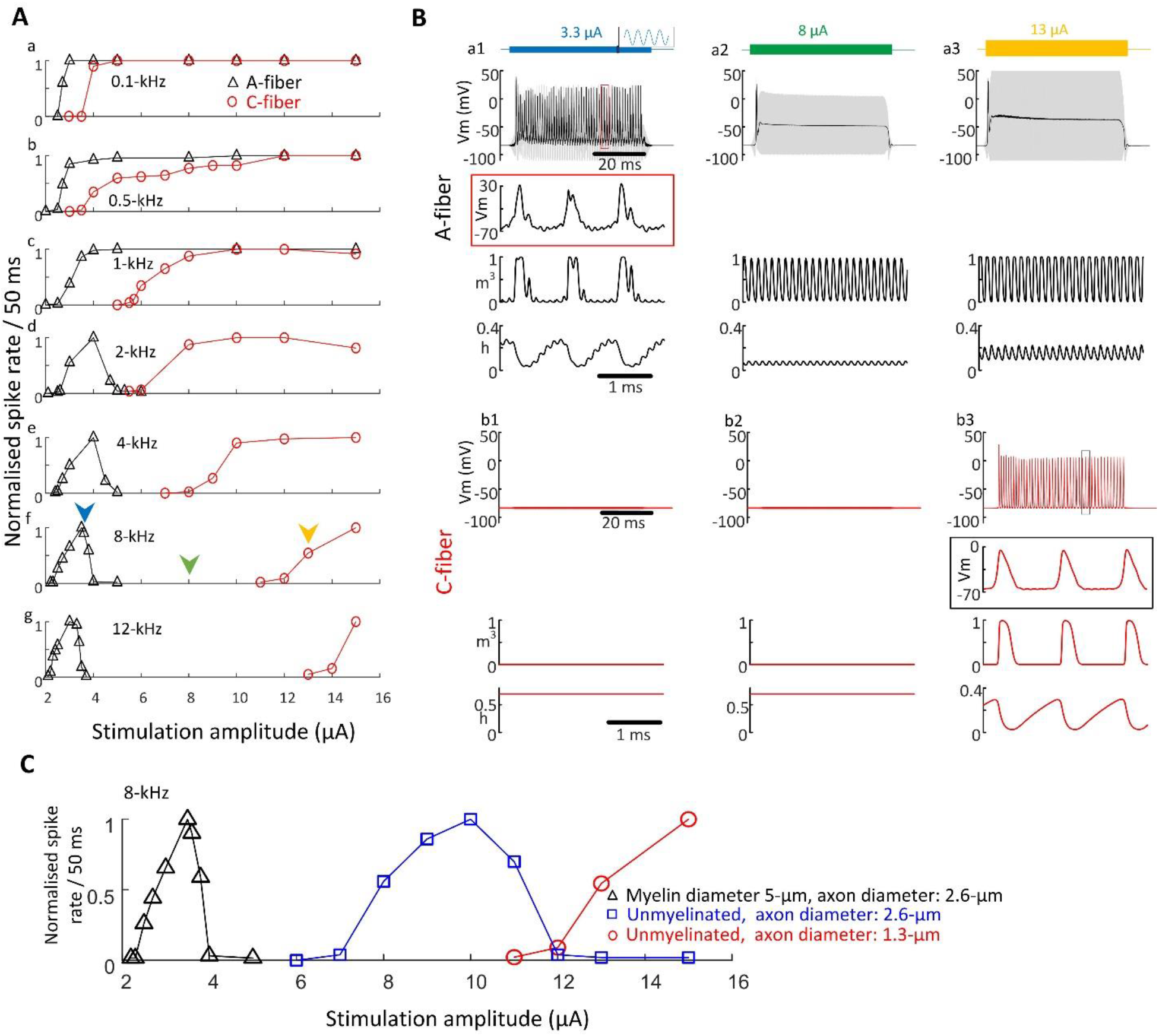
Differential effects of KES on large and small fibers can be explained in computer simulations by how sodium channel responses to stimuli are shaped by axonal size and myelination. Anatomically and physiologically realistic neuro-electric models were implemented to simulate the responses of single fibers to Khz electrical stimuli. (A) Simulated normalized spike rate elicited in the myelinated (A-type) and un-myelinated (C-type) fiber models using 0.1-kHz, 0.5-kHz, 1-kHz, 2-kHz, 4-kHz, 8-kHz and 12-kHz sinusoidal KES, across multiple stimulus intensities. In panel (f), arrowheads point to the 3 stimulus intensities used to compile traces in (B). (B) Examples of membrane voltage (Vm) trajectories (black trace) and stimulus artifacts (superimposed grey trace), along with steady-state activation (m^3^) and inactivation (h) gating variables of sodium current in neurites below the electrode, during 8-kHz KES. Traces are shown for A-fibers (top “a” panels) and C-fibers (bottom “b” panels), at 3 stimulus intensities (3.3μA, left column panels a1 and b1; 8μA, middle column panels a2 and b2; 13μA, right column panels a3 and b3). In panels a1 and b3, short snippets of Vm traces, and the corresponding gating variable time-courses, are shown magnified. (C) Normalized spike rate elicited in the myelinated A-fiber (black), un-myelinated C-fiber (red), and un-myelinated fiber which has the identical axonal diameter with A fiber (blue) by 8-kHz sinusoidal KES

## Discussion

Despite the physiological and translational significance of small, unmyelinated vagal C-afferents, their selective activation using electrical vagus stimulation has not been achieved to date. In this study, we show that intermittent kHz-frequency electrical stimulation (KES) can preferentially engage C-afferents over larger fibers in the vagus and activate C-afferent-associated sensory neurons over motor vagal neurons in the brainstem. This was true in both mice and rats, suggesting a mechanism that is not species-specific.

### KES activates vagal neurons

Our finding that KES of peripheral nerve fibers leads to activation of fiber-associated neurons is surprising, in light of several past studies that have demonstrated KES-elicited block of nerve conduction in general (30) (35, 38) and of vagal C-afferents in particular (35). KES of the sub-diaphragmatic vagus is used clinically in the treatment of obesity(43), putatively acting by blocking conduction in gut-innervating vagal sensory neurons that signal satiety and/or in vagal efferent fibers involved in the control of gastrointestinal fluid release and motility(44). However, our finding suggests that part of the effect of KES in obesity may be mediated by activation of the vagus, as suggested recently(3), rather than nerve block(45).

In our study, we documented neuronal activation by quantifying c-Fos immunoreactivity in neurons in the NTS, a brainstem region that directly receives input from vagal sensory neurons bearing C-afferents, and the DMV, from which much of motor vagal signaling is communicated to the periphery via cholinergic vagal efferents(34). We found that 30 minutes of intermittent KES resulted in increased expression of c-Fos protein in both those regions, at lower but still comparable levels to those produced by “standard” 30 Hz vagus nerve stimulation (Figure 1B-D). Sampling of tissue in our study occurred 1.5 hours after the end of KES (Figure 1A), a time point within the window for strong c-Fos protein expression following neuronal activation(46), suggesting that these c-Fos expression levels likely represent peak neuronal responses to acute nerve stimulation. Compared to 30 Hz VNS, KES elicited a relatively stronger response in NTS than in DMV neurons (Figure 1C-E, Supple. Figure S1D), indicating preferential stimulus engagement of afferent vagal fibers, of which the vast majority are C-type, over efferent fibers, mostly of Aα and B-type(47). Mechanical stimulation of the nerve during acute placement of vagus electrode had a moderate effect on c-Fos expression in NTS, and no effect in DMV (34) (Figure 1B-D). c-Fos expression cannot resolve cells acutely inhibited by a stimulus, therefore the degree to which KES or 30 Hz VNS results in inhibition of neurons in the brainstem is unknown.

### KES alters vagal fiber activity

Direct recordings of ongoing fiber activity during KES were not available to us, as intracellular recordings from single axons are not feasible(48) and extracellular recordings are obscured by the electrical artifact during KES. Therefore, to demonstrate that KES preferentially activates C-afferents over other fiber types in the vagus, we relied on using physiological responses to fiber engagement from some of the end-organs they innervate as proxies for asynchronous fiber activity, e.g. (35, 37, 49, 50, 51). Stimulus-elicited changes in breathing interval (BI) were used to estimate C-afferent activity, since engagement of C-fibers, either optogenetically (Figure 2B-b)(52) or electrically(37) is associated with alterations in the breathing pattern. We found that KES elicits changes in BI and simultaneously minimizes effects on heart rate, an index of B-fiber activation, and on laryngeal EMG, an index of A-fiber activation (Figure 2C-E; Suppl. Video 1D-F; Suppl. Figure S3-S5)(37). When HR changes occur, especially at very high KES intensities (Figure 2D-a, Figure S3A-b), those could be due to C-afferent-mediated vago-vagal reflexes, with slow-onset and less pronounced cardio-inhibition (e.g. Figure 2B-b), as opposed to the fast-onset, intense HR drop seen with direct B-fiber activation(21).

### KES alters vagal fiber excitability

Measurement of physiological responses to neurostimulation is necessary but not sufficient to establish nerve conduction block by KES, nor is nerve block necessarily the cause of a given physiological effect, as this could be achieved through other mechanisms(36), such as facilitation rather than block(53). For the same reasons, physiological responses to KES indicating fiber activation need to be corroborated by direct measurement of changes in fiber excitability, by recording nerve compound action potentials in response to single “probing” pulses (eCAPs) (35, 38, 54). eCAPs in response to probing stimuli before, during and after KES are consistent with sequential conduction block as stimulus intensity increases (Figure 2D and Suppl. Figure S5), which starts with A-fibers, then encompasses B-fibers and finally affects C-fibers (Figure 2D-E), as described before(38, 55). This “intensity window”, within which excitability of A- and B-fibers is almost completely blocked with C-fiber excitability only partially suppressed, provides the basis for the C-afferent selectivity of KES (Figure 2E). It is important to note that asynchronous activity and excitability are 2 different measures of the state of nerve fibers during KES(56): A- and B-fibers have both minimal activity and excitability, whereas C-afferent have higher activity but are also partially suppressed, with regard to their excitability, compared to baseline. Indeed, C-afferent-associated BI responses reach a maximum at intensities producing only half-maximal block of C-fibers in eCAPs (Figure 2C-E), suggesting that action potentials in C-afferents are still elicited by KES trains, even if excitability of those fibers is partially suppressed. This mechanism is consistent with our finding that C-afferent-associated NTS neurons are preferentially activated against Aα/B-fiber-associated DMV neurons, but at sub-maximal levels compared to NTS neuronal activation by non-selective 30 Hz VNS (Figure 1). Changes in fiber excitability occur very fast after the onset of KES and return to pre-stimulation level after the end of KES trains, suggesting that 10-second long trains have a tightly controlled, temporary effect with no damage to fibers(38) (Suppl. Figure S5). Slowing in conduction velocity, more prominent in slow fibers, is observed in our experiments (Figure 2D and Suppl. Figure S5), consistent with previous reports(38).

### Optimization of KES parameters for C-afferents

The effects of KES on vagal fibers are stimulus intensity- and frequency-dependent and non-monotonic (Figure 2C and 2E), and parameters associated with A/B-fiber block and C-afferent activation are different between subjects and species, underlining the need for a “personalized” optimization procedure. In our study, we used selectivity indices (SIs) to normalize and compare responses and determine optimal KES parameters, across subjects (Figures 1E and 3D). These indices account for the fact that the effects of electrical stimulation depend on the nerve-electrode interface and the underlying nerve anatomy(57), both of which are highly variable, and that practically any set of nerve stimulation parameters engages multiple fiber populations, in different degrees, e.g. (40). The nerve potential measurements required for CAP SIs are surgically challenging and often noisy, and the IHC procedures required for neuronal c-Fos SIs are not feasible in real-time. In contrast, compiling SIs from noninvasive physiological responses that represent fiber engagement can be implemented in real-time, and is practical and feasible in experimental animals and human subjects(37). In our study, physiological SIs compiled from A-, B- or C-fiber selective stimuli are strongly correlated with CAP SIs compiled from the same stimuli (Figure 3, Suppl. Figure S6), suggesting that they are good indicators of selective fiber engagement. Such indices could be programed into research or bedside systems for automatic calibration and optimization of VNS therapies targeting different fiber populations in individual patients. For example, B-fiber indices may facilitate calibration of VNS in treating heart failure(58) or cardiac arrhythmias(59), and C-fiber indices in application of VNS in inflammatory(12, 60) or metabolic disorders(61).

In our study, the duration of KES trains, as well the intervals between trains, are likely to have contributed to the C-afferent selective effect. KES trains produce transient fiber excitation followed by longer-lasting inhibition of nerve fibers(38). The intermittent, rather than continuous, time course of KES trains in our study may be responsible for the sustained activation. By controlling the duty cycle, the ON/OFF epochs of KES, the balance between brief excitation and longer-lasting inhibition can possibly be leveraged and further increase the level of C-afferent activation and/or selectivity. Optimizing the duty cycle is important for another reason: a small duty cycle (short ON epochs) might lead to limited selectivity, as a minimum duration is required to block large fibers, and a large duty cycle might end up blocking both large and small fibers.

### A cell membrane mechanism for C-afferent selectivity of KES

We used an anatomically-accurate, biophysically-plausible computational model of axons to study the effects of KES in different regimes, from subthreshold facilitation to suprathreshold block(56, 62), and to gain insight into possible membrane-level mechanisms that could contribute to the differential fiber responses to KES. Our simulations suggest that large, myelinated and small, unmyelinated fibers undergo both activation and block, and that selectivity for one or the other can be attained by modulating stimulus frequency and intensity (Figure 4A), in agreement with our experimental findings. In the model, parameters related to voltage-gated channel kinetics and ion channel distributions were kept the same for large and small fibers; those fibers differed only with regard to their physical properties, i.e. axonal size and presence of myelin(63-65). The activation function of fibers increases with larger axonal radius(66). The decreased axial resistance of larger fibers also leads to greater electrical connection between adjacent axonal segments, preventing the intracellular potential from ‘floating’ with changes in extracellular potential and rendering the axon more responsive to gradients in the extracellular potential. Larger fibers are more strongly affected by KES-induced depolarization and therefore exhibit lower activation and blocking threshold. The effect of myelin does not qualitatively alter but does magnify this effect by significantly increasing the internodal distance, thereby increasing the voltage gradient on each node(66)(Figure 4C). Therefore, morphological differences between fibers with otherwise identical ionic channels can lead to different activation and blocking thresholds, potentially explaining part of the intensity-dependency of KES. he dominant role of sodium channels during KES has been discussed previously(48, 55), as has the contribution of other ionic channels, including delayed rectifier potassium and T-type low-voltage-activated calcium channels(67). Given the limited data on the intrinsic ionic diversity of functionally-distinct vagal fibers(68), our study focused on how properties of “standard”, universally present, sodium channels can shape fiber responses, to gain insight into a minimal set of mechanisms that could explain part of the experimental effect.

The frequency-dependency and fiber-selectivity of KES activation and block can further be explained by the fiber-specific refractory period(69, 70) and passive time constant(56, 71). At subthreshold intensities, for a given fiber type, membrane voltage modulations occurring faster than the membrane time constant, result in charge accumulation, which can further facilitate spiking(56, 72) (Figure 4B-a1 and b3). On the other hand, at suprathreshold intensities, sodium channel inactivation induced by tonic membrane depolarization to consecutive stimuli within refractory period is likely the major mechanism underlying action potential block (Figure 4B-a2 and a3)(73, 74). Intensities that are supra-threshold for large fibers promote conduction block (Figure 4B-a2 and a3), while at the same still being sub-threshold for small C-fibers, facilitating asynchronous C-afferent activity (Figure 4B-b3). By combining the frequency- and intensity-dependencies of KES, the resulting block and activation windows for different fiber types can be leveraged for fiber-selective stimulation (Figure 4A-e through g). The nerve-specific properties may also explain our finding that, during KES, C-fiber eCAPs start to decline at lower intensities than those at which the respective physiological response, i.e. breathing changes, reaches its maximum (yellow bars in Figure 2C and 2E). As KES intensity increases, more asynchronous action potentials are elicited from C-fibers, thereby leading to greater changes in breathing. However, that might also transiently bring more C-fibers into their absolute refractory period, even for a very short time, so fewer C-fibers will be able to generate action potentials in response to the probing stimulus, thus leading to smaller C-fiber eCAPs. It has recently been reported that neural block threshold increases monotonically with increasing frequency between 10 and 300 kHz, when KES is symmetrical(24, 67). Even though we used symmetric KES, in both experimental and modeling studies, we found non-monotonically varying neural block thresholds for A- and B- fibers (Figure 4). A possible reason of this discrepancy could be the different criteria of neural block: block of spike propagation along the axon, adopted in(73), vs. block of the node under the stimulation electrode, adopted in our study. Importantly, a recent study that looked at KES for somatic blocking, in which the stimulation electrode is applied on a cell body, found a non-monotonic neural block threshold with increasing frequency (< 6kHz).

## Methods

### Animal preparation, anesthesia, physiological instrumentation

Forty-two adult male Sprague Dawley rats (age 2-5 months and weight between 300-550 gm) and eleven male C57BL/6 mice (2-4 months and weight between 25-30 gm) were used in the study under the approval of the Institutional Animal Care and Use Committee at The Feinstein Institutes for Medical Research. Rodents were anaesthetized using isoflurane (induction at 4% and maintenance at 1.5-2%) and medical oxygen; anesthesia was maintained throughout the experiment. Body temperature was measured with a rectal probe and maintained between 36.5-37.5°C using a heating pad (78914731, Patterson Scientific) connected to a warm water recirculator (TP-700 T, Stryker). ECG (Figure 2A-b) was recorded by using 3-lead needle electrodes subcutaneously on the limbs and amplified using a commercial octal bio-amplifier (FE238, ADI). Breathing was monitored by using a temperature probe placed outside of the nostrils along with a bridge amplifier (FE221, ADI); the probe reported changes in air temperature during breathing movements: drop in temperature during inhalation, and rise during exhalation (Figure 2A-c). All physiological signals were first digitized and then acquired at 1-kHz (PowerLab 16/35, ADI) and visualized on LabChart v8 (all from ADInstruments Inc).

### Surgical preparation and vagus electrode placement

To expose the cervical vagus nerve (cVN) in the rat model, a midline 3 cm skin incision was given on the neck. Salivary glands were separated, and muscles were retracted to reach the carotid bundle. Under a dissecting microscope, the right cVN was isolated first at the caudal end of nerve and then at rostral end of nerve. The middle portion, between the two isolated sites was left intact within carotid bundle to minimize the extent of surgical manipulation and trauma to the nerve. After isolation of the nerve, a pair of custom-made, tripolar cuff electrodes was placed on the caudal and rostral sites relative to omohyoid muscle (Figure 2A-a). The cuff electrodes were made using a polyimide substrate and sputter-deposited iridium oxide contacts for low electrode impedances and stable stimulation characteristics(75-77). Electrode contacts had dimensions of 1418×167 µm^2^ with an edge-to-edge spacing of 728 µm and center-to-center spacing of 895 µm. Typical individual electrode impedances in saline ranged from 0.5 to 1.5 kΩ. The distance between the stimulating electrode (center contact of tripolar cuff) to the most proximal recording electrode on the nerve was measured roughly 5 to 6 mm. Silicone elastomer (Kwiksil, World Precision Instruments) was placed around the cuff to minimize current leakage during stimulation. In the mouse model, all surgical procedures were identical except the left cVN was targeted. In addition, for direct laryngeal muscle measurement, the thyroid cartilage was exposed by separating the sternohyoid muscle at the midline using blunt dissection. Using a 29G insulin syringe, a shallow slit was made in the thyroid cartilage just lateral and inferior to the laryngeal prominence. With the needle bevel facing up, the two PTFE-coated platinum-iridium wires were carefully inserted into the underlying laryngeal muscles through the slit guided by the syringe needle.

### Vagus nerve recording and stimulation

Neural activity from each contact on the recording electrode was amplified, digitized (30KS/s, 16bit resolution) and filtered (60-Hz notch), using a 32-channel RHS2000 stim/record headstage and 128ch Stimulation/Recording controller (Intan Technologies); recordings were single-ended, relative to a ground lead placed in the salivary gland. Nerve stimulation was delivered in constant current mode as trains of pulses using an STG4008 stimulus generator (Multichannel Systems). For all experiment related to waveform manipulation, the stimulation waveforms were composed of monophasic pulse with varying pulse width, intensity, polarity, and shape. Monophasic pulses were used here to yield lower threshold and simpler stimulus artifact shape. Even though pulses were not charge-balanced, it is unlikely that during these acute experiments with low pulsing frequency, significant charge build-up occurred. In particular, fully randomized single pulse with 30-s on and 10-s off at 1Hz were used to access the neural response, whereas stimulus trains of 10-s durations with identical type of pulse at 30Hz were randomly delivered to evoked discernible physiological response. For experiments related to frequency manipulation, all the stimuli were delivered in biphasic form except for probing pulse, to maintain the charge balancing across the neural interface and minimize the neural injury. All the stimuli were constructed as trains with consistent 10-s duration but with varying frequency, pulse width, and intensity, and randomly delivered. The stimulation configuration was tripolar (cathode-center or cathode-corner) as it outperforms in terms of protection of current leakage for all experiments. There were at least 15-s long pauses between successive trains to ensure that physiological measurements had reached a steady state before a new train was delivered. For optogenetic stimulation, ChAT-IRES-Cre (#006410), Vglut2-IRES-Cre (#016963), and Ai32 ChR2-eYFP (#024109) mice were purchased from The Jackson Laboratory and crossed to produce ChAT-ChR2-eYFP and Vglut2-ChR2-eYFP mice. 8- to 16-week-old mice were anesthetized and the vagus nerve exposed as described earlier. A custom-made optical cuff, consisting of a blue LED light source (XLAMP XQ-E, Cree LED) in a molded silicone enclosure (Nusil MED-4211), was placed on the left cervical vagus nerve and connected to a stimulus generator (Multichannel Systems). Optogenetic stimulation was delivered using 10-sec stimulus trains of 10 ms pulse width and 30 Hz frequency while recording heart rate and breathing rate responses. Optical stimulus intensity was varied by gradually increasing the voltage driving the LED, with threshold of visual perception around 2.3V.

In all experiments with neural recording, we initially determined the “neural threshold” or threshold (T) as the lowest stimulus intensity for a 100-µs duration pulses that evoked a discernible evoked potential at the recording electrode. The physiological threshold (PT), which evoked visible (5-10%) heart rate/respiratory change (usually 3 or 4×NT), was used in experiment when no neural signals were recorded and for all KES experiments. The details of each experiment can be found in Table S1 and S2. Specifically, to access the neural activity in response to the KES with one stimulation cuff, we designed the waveform, which is combined with KES with low frequency, 1Hz probing pulse, as the low frequency probing pulse does not contribute significantly to physiological effect(37). For each probing pulse, a 30-ms window (5ms before, 25ms after the onset) is opened in the 10-s KES train, to improve the signal-to-noise ratio for further evoked neural signal processing in next section.

### Identification and analysis of neural and EMG signals

Raw nerve signal traces from both electrodes were filtered using a 1Hz high-pass filter to remove the DC component. Stimulus-evoked compound action potentials (eCAPs) elicited from individual pulses or from trains of pulses, were extracted, by averaging individual sweeps of nerve recording traces around the onset of pulses (waveform manipulation experiments) or probing pulse (frequency manipulation experiments). A custom-made buffer amplifier was used to record the induced voltage on the electrode during stimulation. Stimulation artifact was suppressed offline for waveform manipulation experiment by a recently proposed method which subtracts the trace of the stimulation electrode voltage from the eCAPs with proper template matching and an edge effect removal algorithm(39). For frequency manipulation, due to the saturation of artifact voltage buffer, same artifact removal algorithm has not been applied.

Given the rough estimation of distance between the recording and stimulation electrodes (5-6 mm), we fine tune the distance in analysis so that the latency windows can align well with the A-, B- and C-fiber prominent peaks with pre-defined conduction velocity ranges for each fiber type (A: 5-120 m/s; B: 2-8 m/s; C: 0.1-0.8 m/s)(78). Figure **1**2A-d shows representative eCAPs, including activity of different fiber types and EMG. Signals from both contacts in the recording electrode, proximal and distal to the stimulating electrode, were collected (solid black and dashed red traces in Figure 2A-d). This allowed to distinguish between neural and EMG signal components. For the given electrode spacing A- and B-fibers had short latencies (< 3 ms, red and green windows), while slower C-fibers occurred at longer latencies (> 6 ms, yellow window)(39). To discriminate C-fiber components from EMG, we reasoned that C-fiber volleys should show a latency difference between the proximal and distal recording contact, spaced apart by a distance of 895 μm, of 1-2 ms, whereas EMG signals should occur simultaneously on both recording contacts(39) (Figure 2A-d, grey window), with time window around 2-6 ms (identified with neuromuscular junction blocking agent in our previous study(37)).

### Analysis of physiological signals

We computed the magnitude of EMG response from respective eCAPs as the peak-to-trough amplitude of the (typically biphasic) response within the EMG window (Figure **1**2A-d, grey window).; that amplitude was then normalized by the maximum EMG amplitude in that subject. Using a custom algorithm, ECG peaks corresponding to the R waves were identified, and heart rate (HR) was computed from R-R intervals. We defined stimulus-induced change in HR (ΔHR) as the difference between the mean HR during a 10-s epoch before the onset of the stimulus train and the mean HR during the stimulus train (“VNS”), divided the mean pre-stimulus HR (Figure **1**2A-b). In recordings from the nasal temperature sensor, we identified peaks (end of expiration) and troughs (end of inspiration). We defined the interval between two successive peaks (or two successive troughs) as breathing interval (BI). We defined the stimulus-elicited change in breathing interval (ΔBI) as the difference between the mean pre-stimulus and the mean during-stimulus BI (Figure 1A-c). For those cases without breath during stimulation period, the breathing interval between last breath of pre-stimulus and first breath post-stimulus was used as mean during-stimulus BI. The measured signals and corresponding derived variables (ECG and ΔHR, and nasal sensor temperature and ΔBI) are shown in Figure **1**2A-b and -c respectively. All the analyses were performed using MATLAB 2016a software (MathWorks, Natik, MA, USA).

### Immunohistochemistry

Rats delivered VNS intermittently for 30 minutes (10 seconds on, 50 seconds off), were deeply anesthetized with isoflurane and transcardially perfused using 250ml of 0.9% saline after 1.5 hrs. The brains were removed immediately and post-fixed for 2 days in 4% paraformaldehyde in 0.1 M PBS. After post fixation, the brainstem was precut by razor blade and sectioned in vibratome with 50µm thickness (VT1200S, Leica, U.S.A). Sections, acquired from anterior-posterior: - 13.4mm - -13.9mm relative to Bregma, were washed with TBS buffer 3 times for 5 minutes and subjected to antigen retrieval using 1X SignalStain® Citrate Unmasking Solution (Cell signaling). The citrate buffer was bought to boiling temperature and added to the sections. The well plate was incubated at 85°C for 10 minutes. The plate was allowed to cool down and the sections were washed with TBST buffer (1X TBS buffer + 0.025% Tween 20). 5% normal donkey serum and 0.3% Triton-X100 in TBS buffer was used as blocking buffer and the sections were blocked for 1 hour at room temp. Sections were first stained with c-Fos and then Choline Acetyltransferase (ChAT) was performed subsequently. c-Fos staining was the done by incubating the sections with the c-Fos antibody (Abcam, ab190289) at 1:1000 dilution in blocking buffer for 3 days placed in shaking incubator at 4°C. Section were rinsed with the TBST buffer 3×5 times and incubated in the donkey anti-rabbit secondary antibody labeled with alexa-fluro 488 (1:500) in blocking buffer for 2 hours at room temperature. Sections were washed 3×5 minutes in TBST buffer and incubated with ChAT antibody (Sigma, AB144) at 1:100 dilutions overnight at 4°C in the blocking buffer. Sections were rinsed the next day with 3×5 minutes in TBS buffer and incubated with anti-goat secondary antibody labelled with alexa fluro 555 (1:200) for 2 hours in room temperature. Sections were rinsed in TBST buffer 3 times and incubated with DAPI 1:1000 dilution in TBS buffer for 1 hour at room temperature. The section was then rinsed two times with TBS buffer and mounted on to Poly-L-lysine coated glass slides. Cover glass was secured on the top of the sections with VECTASHIELD® PLUS Antifade Mounting Medium (Vector labs, H-1900). Sections from naïve, sham and different VNS treatment group were processed in parallel.

DAPI, c-Fos and ChAT expression in the nucleus tractus solitarius (NTS) and dorsal motor nucleus of the vagus nerve (DMV) was captured using all-in-one fluorescence microscope under 20X objective (BZ-X800, Keyence, U.S.A.), with region of interests (medial-lateral: ±2mm relative to mid sagittal plane, dorsal-ventral: 7.2-8.2mm relative to brain vertex). After processed by commercial imaging stitching software (BZ-X800 Analyzer), the orientation of the stitched images was first adjusted for cross-animal comparison and consistency. To correct non-uniform illumination and attenuate background, the original image was subtracted from background approximation image estimated from morphological opening method. ROIs for NTS and DMV regions were marked by anatomical expert though DAPI and ChAT staining. Cells expressing c-Fos were then counted bilaterally in 3-4 sections/brain using thresholding, watershed separation, and automatic particle counting tools in ImageJ. All the image post-processing techniques were performed using ImageJ or MATLAB 2016a.

### CAP, physiological, and neuronal-c-Fos selectivity indices

To evaluate the fiber selective performance of tested stimulation parameters, for different types of fibers, we defined CAP selectivity indices (CSIs) which aim to maximize target and minimize off-target fiber activity. The CSIs for A-, B-, C-fibers are:

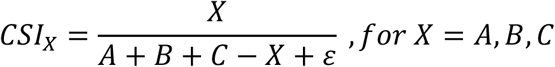

where A, B, C are normalized evoked fiber activity amplitude within each animal, and a small constant ε is used to prevent the overflow due to the extremely small fiber activity in the denominator.

Similar to CSIs, based on an existing relationship/models between fiber activity and physiological response: A-fiber to evoked EMG, B-fiber to HR, and C-fiber to BI (Figure S2)(37), we defined physiological selective indices (PSIs) which aim to maximize desired and minimized non-desired physiological effects corresponding to different type of fiber activation. The PSI for A-, B-, C-fibers defined as:

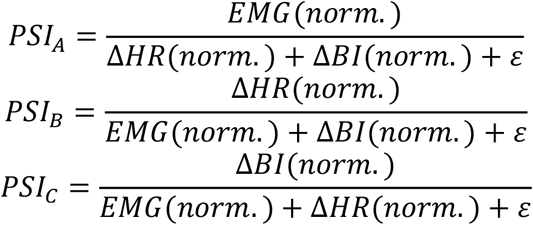

where EMG(norm.), ΔHR(norm.), and ΔBI(norm.) are normalized physiological responses within each animal, and a small constant ε is used to prevent the overflow, same as CSI. To quantify the performance of PSIs using different stimulation parameters across animals, a 1st order Gaussian model was computed to capture the relationship between computed PSIs and different stimulation intensities.

Finally, to quantify the immunohistochemistry results in brainstem regions, we further defined neuronal-c-Fos^+^ selectivity index (NcSI) for sensory neurons (mostly related to C-fibers), which is:

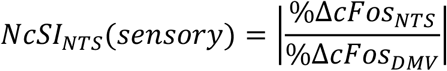

where Δc-Fos in NTS and DMV are normalized with respect to the average number of expressed neurons in corresponding regions in sham group animals.

### Computational model of vagal fibers

All simulations were implemented in COMSOL Multiphysics v. 5.4 (COMSOL Inc., Burlington, MA). Our nerve fiber model structure and parameters were adapted from the McIntyre, Richardson and Grill (MRG)(79) and the Schwarz, Reid and Bostock (SRB) models(80). Two major nerve fiber subtypes were simulated; myelinated A fiber and unmyelinated C fibers. Ion channels are modelled on the nodes of Ranvier, based on the formulations of the SRB model according to:

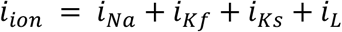

As shown in Figure S14A, the extracellular environment was modelled by a 1000-µm long, 40-µm diameter cylinder surrounding the 1D nerve fiber(81). Two 50-µm electrodes (50 µm apart) were placed on the surface of the cylinder with the electrode edges forming a 60° angle with the nerve fiber. The first electrode was the cathode and the second was designated as ground.

The stimulus waveform included a wide range of frequencies ranging from 0.1-kHz to 12-kHz sinusoid KES, with a duration of 50 ms. A no-flux (i.e insulating) boundary condition was implemented for *V*_*i*_ and *V*_*e*_ at the ends of the fiber. The mesh for the myelinated fibers was set to a total of 20 elements for each myelin segment and a size of 0.5 µm for each node segment. The mesh for nonmyelinated fibers was set to a total of 20 elements for each fiber segment, defined as being the same length as the myelin segments of the myelinated fibers. The length of the nodes was set to 1 µm in all myelinated fibers(79). The length of the myelin compartment was also modelled as a function of the myelin diameter(79, 82). The node and myelin diameters used in the model were estimated based on histological data from rat cervical nerves(82). The model’s predictive ability was validated by in vivo compound nerve action potential recordings from the same animals(82).

Node and myelin structures in the model fibers were characterized by different partial differential equations (PDEs). Myelin was approximated by a distributed resistance in parallel with a capacitance (Figure S14B). We approximated the MRG double cable structure by a single-cable model of the vagus nerve to reduce the computational complexity. The membrane dynamics at the node follows SRB formulations. Models for all fiber types shared ion channel parameters but had fiber-specific physical parameters. All model parameters are listed in Table S3.

The extracellular potential distribution *V*_*e*_ was calculated using:

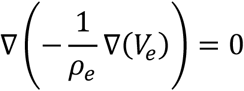

where *ρ*_*e*_ is the extracellular resistivity. The intracellular potential *V*_*i*_ was calculated separately for the myelin and node compartments:

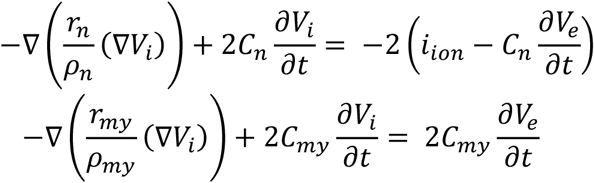

where *r*_*n*_ and *r*_*my*_ are the nodal and myelin radius respectively. Membrane potential *V*_*m*_ was determined from the difference between the intracellular and extracellular potentials.

### Statistics

Analysis of Covariance (ANCOVA) was used to compare the neural responses (A-, B-, C-), physiological responses (EMG, HR, BI), and proposed CSIs and PSIs for different stimulus manipulations (categorical independent variable) and intensity (continuous independent variable). Linear regression was used to compare the same stimulus parameter with different intensity. One-way analysis of variance (ANOVA) and Tukey’s post-hoc tests were used to compare the histological results in brainstem, and two sample t-test was used for corresponding NcSI. Comparison were deemed statistically significant for *p<0*.*05* for all analyses. All statistical analyses were performed on MATLAB (Mathworks).

## Author contributions

YCC conceived and designed experiments, performed experiments, analyzed and interpreted experimental results, and wrote the paper. UA conceived and performed rat experiments. NJ and YCW designed and performed anatomy experiments. IM, MG, and AA designed and performed experiments in mice. QL, TG, and SD designed and performed the computational stimulation, and wrote the paper. AG performed rat experiments. AD and JA performed the histological analysis. SD, TDC, and YAA critically reviewed the paper. SZ conceived and designed experiments, secured funding, analyzed and interpreted experimental results and wrote the paper.

## Data and material availability

All data that support the findings of this study are available from the corresponding author upon reasonable request.

