## Supplemental Materials for "Intermittent KHz-frequency electrical stimulation selectively engages small unmyelinated vagal afferents"

### Supplementary Materials

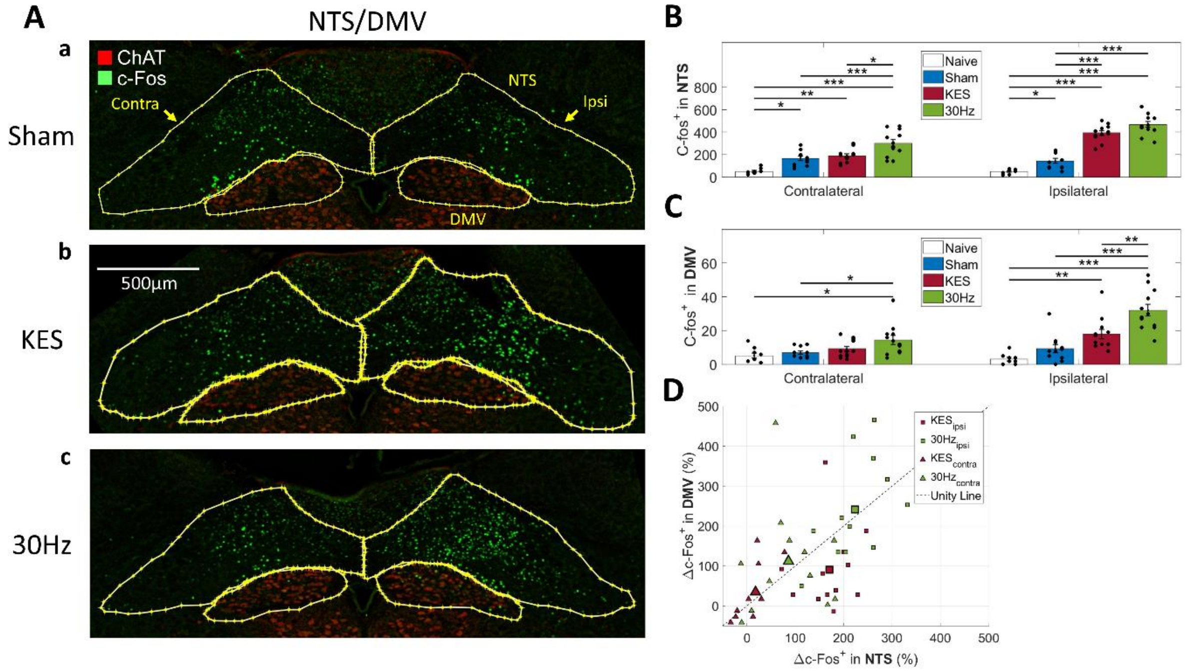


**Figure S1: c-Fos expression in sensory and motor vagal brainstem regions after 30Hz and 8 kHz VNS, in the rat model.**

(A) Representative immunohistochemistry images of sections across ipsilateral and contralateral, to VNS, sensory and motor brainstem regions (yellow contours): nucleus tractus solitaries (NTS) and dorsal motor nucleus of the vagus (DMV) (anterior-posterior: -13.4mm - 13.9µm relative to Breagma, medial-lateral: ±2mm relative to mid sagittal plane, dorsal-ventral: 7.2-8.2mm relative to brain vertex), each stained for c-Fos (green) and ChAT (red). (a) After sham stimulation (electrode placed on nerve, no stimulation delivers). (b) After 30 min of intermittent 8 kHZ VNS (KES, 10-s ON and 50-ss OFF, 30 cycles) (c) After 30 min of intermittent 30 Hz VNS, of the same cyclic settings.

(B) c-Fos$^{+}$ cell numbers (mean±SE) in contralateral and ipsilateral NTS, in different group of animals: naïve (white), sham stimulation (blue), KES(red), 30 Hz VNS (green). Each group consisted of 3 animals and each animal contributed 3-4 sections from which cell numbers were calculated. Statistical comparisons between groups used one-way ANOVA and Tuckey's post-hoc tests (*$p<0.05$, **$p<0.005$, ***$p<0.0005$).

(C) Same as (B), but for DMV.

(D) Normalized c-Fos$^{+}$ cell numbers (% difference in c-Fos+ cell numbers with respect to average cell numbers in sham group) counted in DMV vs. those counted in NTS (ipsilateral: square dots, contralateral: triangle dots), from all sections from 30 Hz VNS group of animals (green dots) and from the KES group of animals (red dots). Mean values shown with black outlined dots.

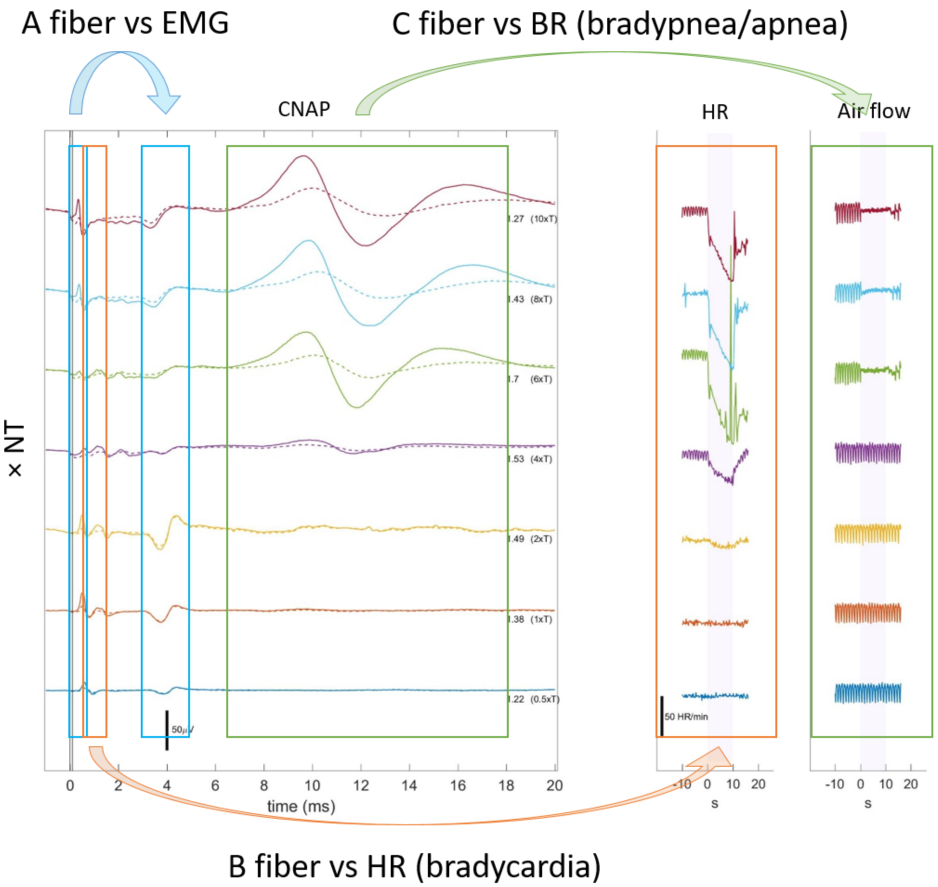


**Figure S2: Electrical activation of different vagal fiber types and their corresponding physiological effects.** A, B, C fiber activities significantly correlate with EMG, HR, respiratory patterns respectively.

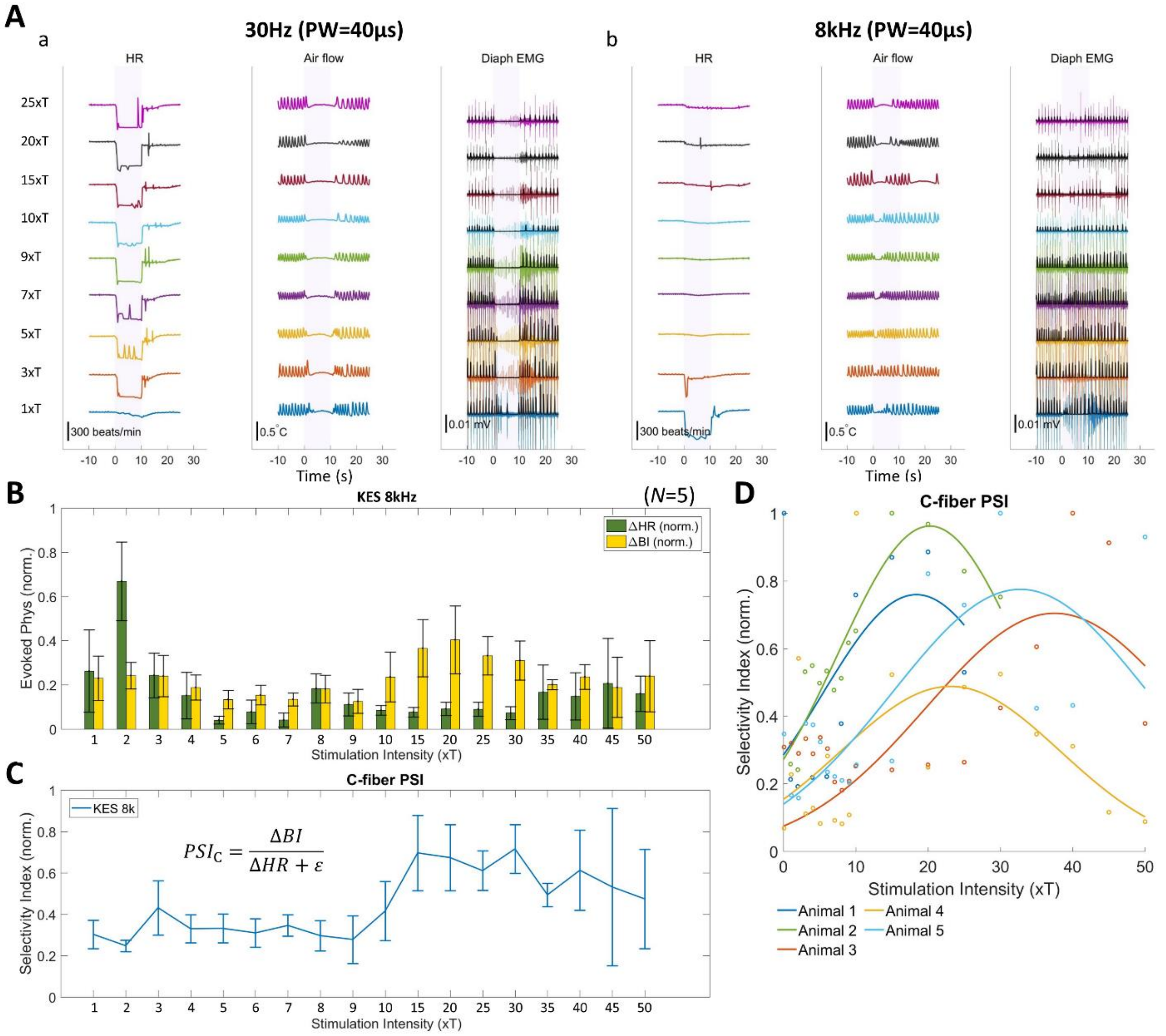


**Figure S3: Elicited physiological effects and resulting physiological selectivity indices (PSI), in response to 8-KHz frequency electrical stimulation of the vagus (KES) of different intensities, in the mouse model.**

(A) Representative ΔHR, ΔBI, and diaphragm EMG (associated with breathing pattern) responses by 30Hz and KES in a single animal, at different stimulus intensities. Stimulus intensities are expressed in units of threshold intensity, defined as the PT of KES (1 to 25×PT).

(B) Mean (±SE, N = 5 animals) of normalized magnitude of physiological responses (ΔHR, green; ΔBI, yellow), elicited by KES of different intensities. (Linear regression, HR: $p=0.156$, BR: $p=0.32$, for intensity).

(C) Mean (+/-SE) values of PSI for C-fibers as function of KES intensity, in 5 animals. (Linear Regression, $p<0.05$ for intensity).

(D) Normalized PSI values for C-fibers cross animals, and Gaussian function is fitted to show the distribution of PSIs evoked by KES stimulus. Average Rmse for the fits: 0.211.

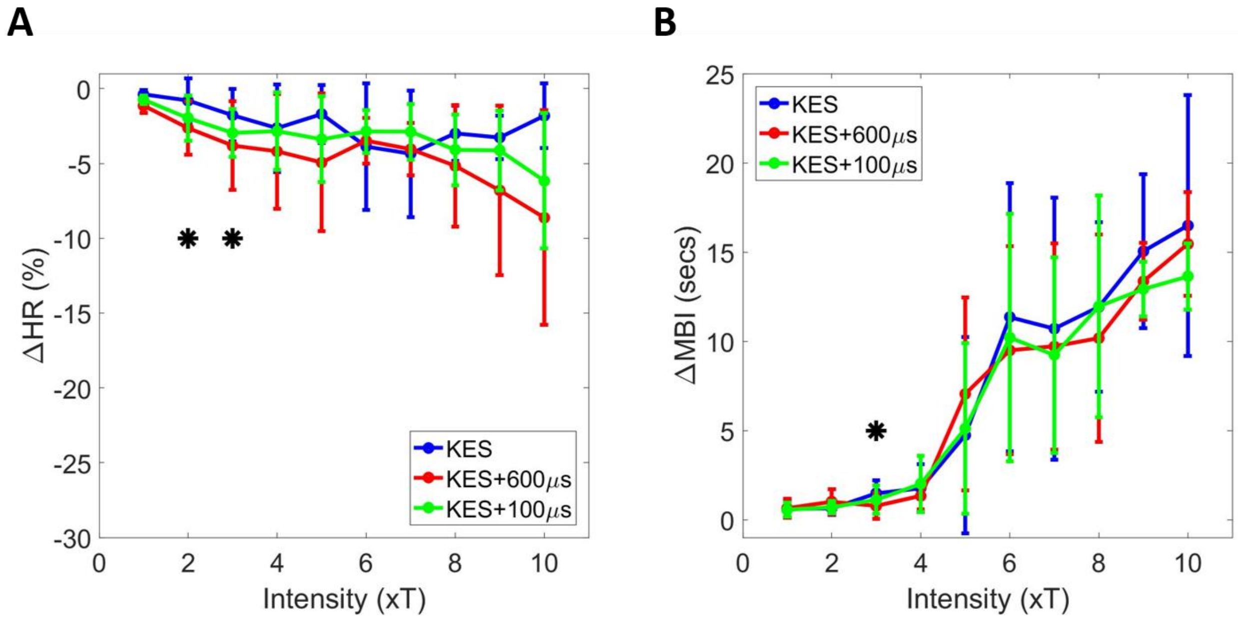


**Figure S4: Comparison of physiological responses evoked by regular KES stimuli and with different type of probing pulses.** (A) The capture curves of ΔHR across animals (N=4) in response to KES (blue), KES with 600µs probing pulse (red), and KES with 100µs (green) probing pulse. The lines and bars represent the average and standard error of the ΔHR respectively. The intensity labelled with star symbol (*) represent the responses evoked by either two type of stimuli showing statistical significance (Pair-wise t-test, $p<0.05$). (B) Same as (B) but for ΔBI.

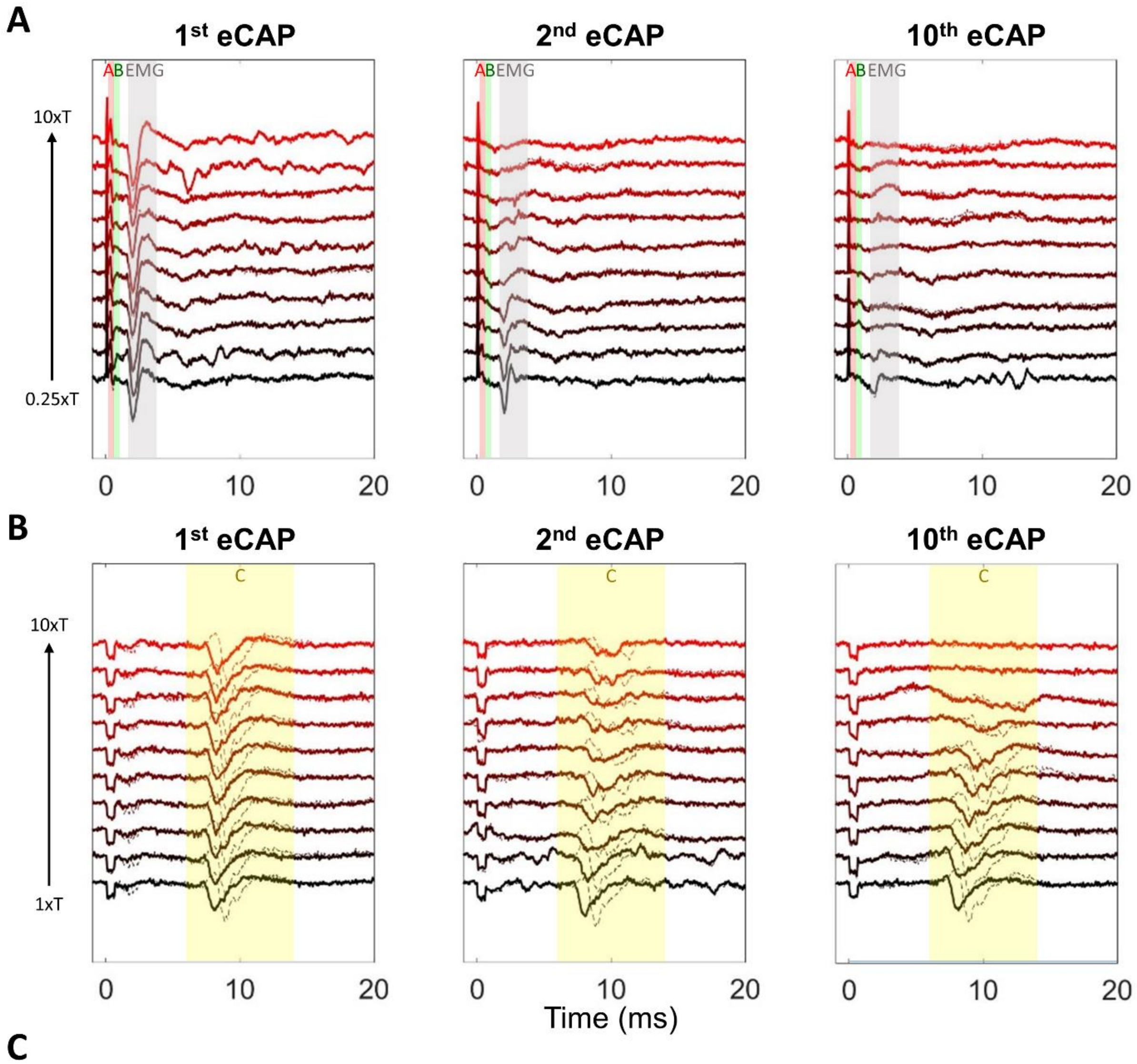


C

| | Comparison | Significance |
|---|---|---|
| A-fiber amp (norm.), 1st eCAP | 0.25xT vs 10xT | P = 0.94 |
| A-fiber amp (norm.), 10th eCAP | 0.25xT vs 10xT | P < 0.05 |
| C-fiber amp (norm.), 1st eCAP | 1xT vs 10xT | P = 0.59 |
| C-fiber amp (norm.), 10th eCAP | 1xT vs 10xT | P < 0.05 |

**Figure S5: Fiber component of eCAPs change over KES stimuli.** (A) Representative 1st (before KES), 2nd (during KES), and 10th (end of KES) eCAPs and evoked EMG elicited by 100μs probing pulse, from low to high KES intensity. The latency windows for the A-, B-fiber and EMG component are shown with red, green, and grey shaded areas, respectively. (B) Same as (B), but for eCAPs elicited by 600μs probing pulse. The latency windows for the C-fiber component is shown with yellow shaded area. (C) Statistical analysis of different KES intensity, for 1st and 10th normalized A-fiber and C-fiber amplitude from eCAPs (Two-sample t-test).

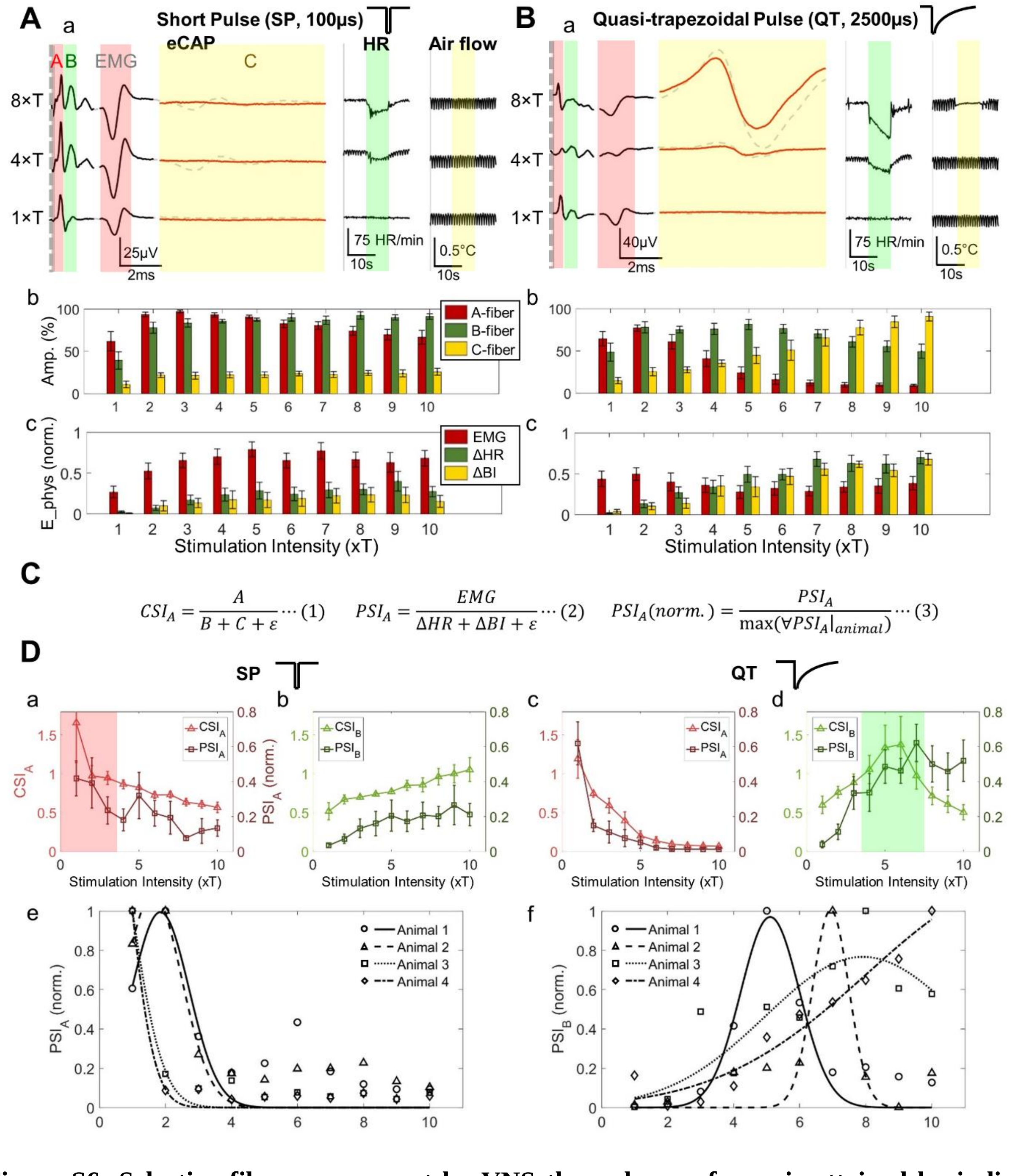


$$CSI_A = \frac{A}{B + C + \varepsilon} \cdots (1) \qquad PSI_A = \frac{EMG}{\Delta HR + \Delta BI + \varepsilon} \cdots (2) \qquad PSI_A(norm.) = \frac{PSI_A}{\max(\forall PSI_A|_{animal})} \cdots (3)$$

**Figure S6: Selective fiber engagement by VNS through waveforms is attained by indices capturing compound action potential (CAP) and physiological responses.**

(A) Responses of trains of square pulses with short pulse width (SP, 30 Hz, 100 µs), known to activate A-fibers. (panel a) Representative eCAP (left panels), HR (middle panels) and breathing responses (right panels) to increasing, from bottom to top, VNS intensities (expressed in units of threshold—the minimum intensity that generates a response). At low intensities, A-fibers and laryngeal EMG increase in parallel. At higher intensities, B-fibers are activated, in parallel with HR changes. No C-fiber (red traces, differential signal from two recording contacts) or breathing responses are elicited with short square pulses. (panel b) Mean (±SE, N = 5 animals) of normalized amplitudes of A-, B-, C-fiber activity (red, green and yellow bars, respectively) evoked by short square pulses at different intensities. (panel c) Mean (±SE, N=8 animals) of normalized magnitudes of physiological responses: EMG, HR (ΔHR) and breathing interval changes (ΔBI) (red, green and yellow bars, respectively) elicited at different intensities.

(B) eCAP and physiological responses for trains of quasi-trapezoidal pulses (30 Hz, 100μs plateau-2500μs falling time) (panel a), known to selectively activate B-fibers[32]. At intermediate intensities, selective B-fiber activation with minimal A-fiber activation (panel b) and the associated ΔHR response with minimal EMG (panel c) are observed. C-fibers are activated at higher intensities, along with the associated ΔBI response (panels a-c).

(C) For a given fiber type, we defined eCAP SI (CSI) as the ratio of eCAP activity of the given fiber, divided by the sum of eCAP activities of the other fibers. Similarly, physiological SI (PSI) was defined as the ratio of the normalized magnitude of the physiological response corresponding to asynchronous activity of the given fiber, divided by the sum of responses for the other fibers.

(D) (panel a) Mean (±SE, N = 5 animals) A-fiber CSI and normalized PSI (N = 8 animals) calculated for short square pulse 30-Hz trains. Both indices are highest at low intensity (red-colored window). (b) B-fiber CSI and PSI for short square pulse trains (ANCOVA, $p<0.05$ for all types of CSI/intensity interaction; $p<0.05$ for all types of PSI/intensity interaction). (c-d) Same as (a-b) but for quasi-trapezoidal (QT) pulse trains (100 μs-plateau followed by a 2500 μs exponentially decaying phase) (ANCOVA, $p<0.05$ for all types of CSI and intensity, and CSI/intensity interaction; $p<0.05$ for all types of PSI, and PSI/intensity interaction). QT pulses of intermediate intensity produce highest B-fiber selectivity (green-colored window). (e) A-fiber PSI values for SP at different intensities and Gaussian fits in representative animals. Average Rmse for fits: 0.133. (f) Same as (e), but B-fiber PSI values fot QT. Average Rmse for fits: 0.159.

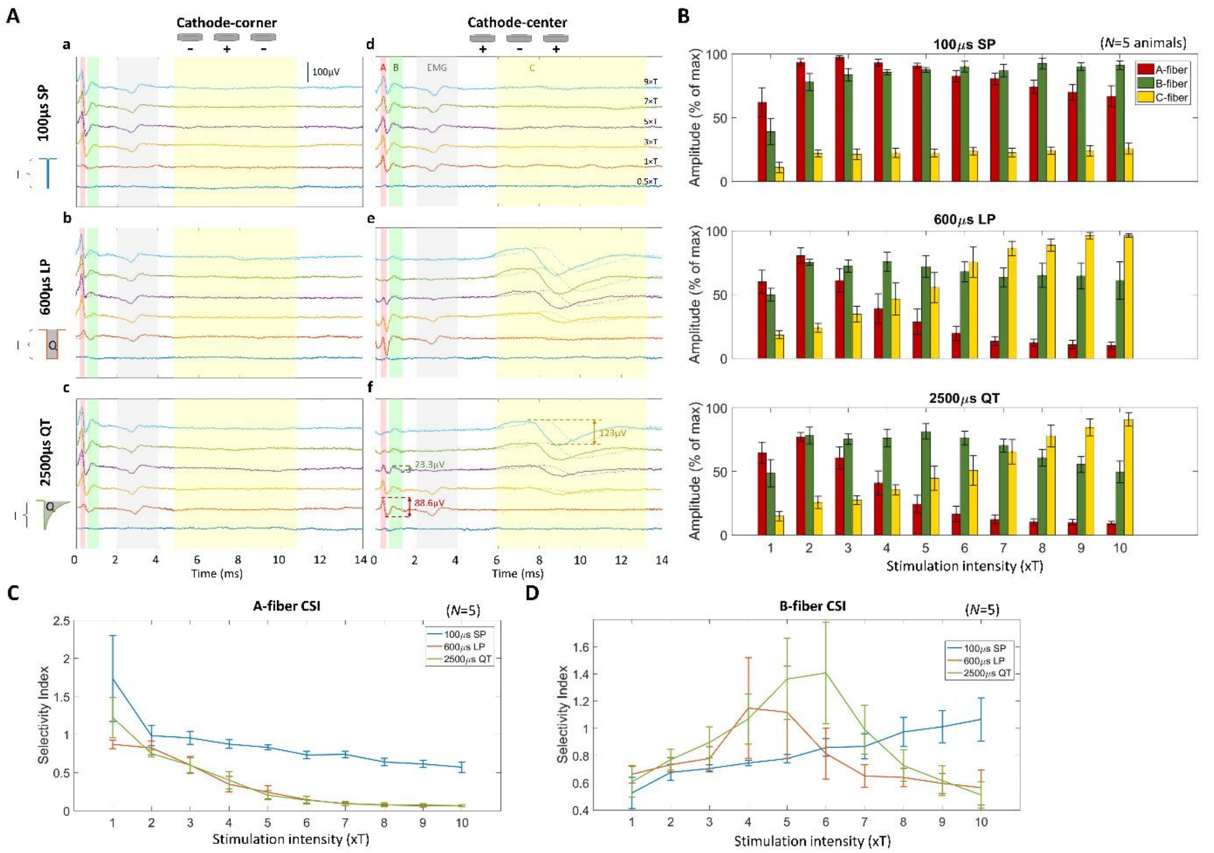

**Figure S7: Stimulus-evoked compound action potentials (eCAPs), fiber activation amplitudes and resulting CAP selectivity indices (CSIs) in response to A- and B-fiber-selective stimulus waveforms, in the rat model.**

(A) Representative eCAPs evoked by different stimulus waveforms: 100 µs square pulse (short square pulse, SP; panel a, d), 600 µs square pulse (long square pulse, LP; panel b, e), and 100 µs-plateau followed by a 2500 µs exponentially decaying pulse (quasi-trapezoidal pulse, QT: panel c, f). Shown are representative eCAP examples in response to cathode-center and cathode-corner polarities and a range of stimulus intensities, from 1 to 10 times thresholds (×T) in a single animal. Traces from both proximal (solid) and distal (dash) contacts of the recording electrode are shown and each trace represents the average of triggers. A-, B-, and C-fiber activation and stimulus-evoked EMG activity, are calculated as peak-to-trough amplitude of eCAP components with different latencies and their overlap with latency windows corresponding to the conduction velocities of different fibers (A-fiber: red; B-fiber: green; C-fiber: yellow; EMG: grey).

(B) Mean (±SE, N = 5 animals) normalized fiber activity evoked by the 3 stimulus waveforms of cathode-center polarity, at different stimulus intensities, in 5 animals. (ANCOVA, *p<0.05* for waveform, across all fibers (A-, B-, C-), *p<0.05* for intensity except of B-fibers (0.21)).

(C) Mean (±SE, N = 5 animals) CSI values for A-fibers, for each of the 3 stimulus waveforms, as a function of stimulus intensity. (ANCOVA, *p<0.05* for waveform and intensity, *p>0.05* for interaction).

(D) Same as (C) but for B fibers. (ANCOVA, *p=0.27* and *p=0.75* for waveform and intensity respectively, *p<0.05* for interaction).

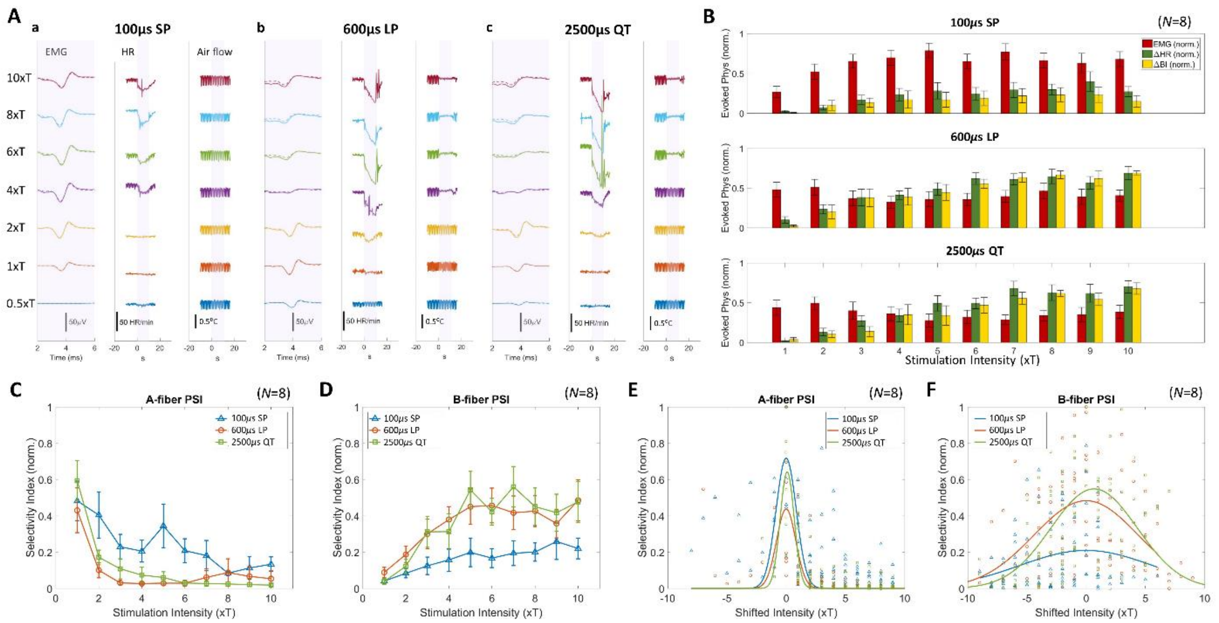


**Figure S8: VNS-elicited changes in physiological parameters, magnitude of physiological effects and resulting physiological selective indices (PSIs) in response to A- and B-fiber selective stimulus waveforms, in the rat model.**

(A) Representative laryngeal EMG (left for each panel), heart rate (HR, middle of each panel), and breathing responses (air flow, right of each panel) elicited by stimulus trains of 3 different waveforms, short pulse (SP), long pulse (LP) and quasi-trapezoidal (QT), for a range of stimulus intensities (0.5 to 10×T).

(B) Mean (±SE, N=8 animals) normalized EMG (red bars), HR (ΔHR, green bars) and breathing interval responses (ΔBI, yellow bars) elicited by the 3 stimulus waveforms, for different stimulus intensities. (ANCOVA, $p<0.05$ for waveform, across all physiological responses (EMG, HR, BI), $p<0.05$ for intensity, across all physiological responses except for EMG (0.9)).

(C) Mean (±SE, N=8 animals) normalized A-fiber PSI values associated with each of the stimulus waveforms (SP: blue, LP: red, QT: green trace), as a function of stimulus intensity. (ANCOVA, $p<0.05$ for waveform and intensity, $p>0.05$ for interaction).

(D) Same as (C) but for B-fibers. (ANCOVA, $p<0.05$ for waveform and intensity, p<0.05 for interaction).

(E) Normalized A-fiber PSI values for each of the 3 waveforms plotted against stimulus intensity, re-aligned to the value associated with the maximum PSO for each animal and waveform, so the intensity shift of 0 corresponds the intensity producing the maximum PSI for each animal. Also shown are 1-D Gaussian function fits to the data for each of the 3 waveforms. Rmse for each fits: 0.235(SP), 0.126(LP), 0.111(QT). (ANCOVA, $p<0.05$ for waveform and absolute intensity shift, $p>0.05$ for interaction).

(F) Same as (E) but for B-fiber PSI values. Rmse for each fits: 0.188(SP), 0.240(LP), 0.254(QT). (ANCOVA, $p<0.05$ for waveform and absolute intensity shift, $p<0.05$ for interaction).

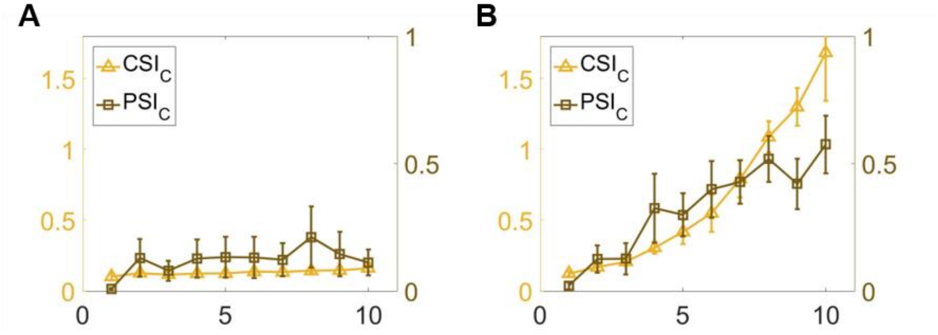


**Figure S9: C-fiber CSI and PSI for short square and quasi-trapezoidal pulse trains.** (A) Mean (±SE, N = 5 animals) C-fiber CSI and normalized PSI (N = 8 animals) calculated for short square pulse 30-Hz trains. (B) Same as (a) but for QT pulse trains.

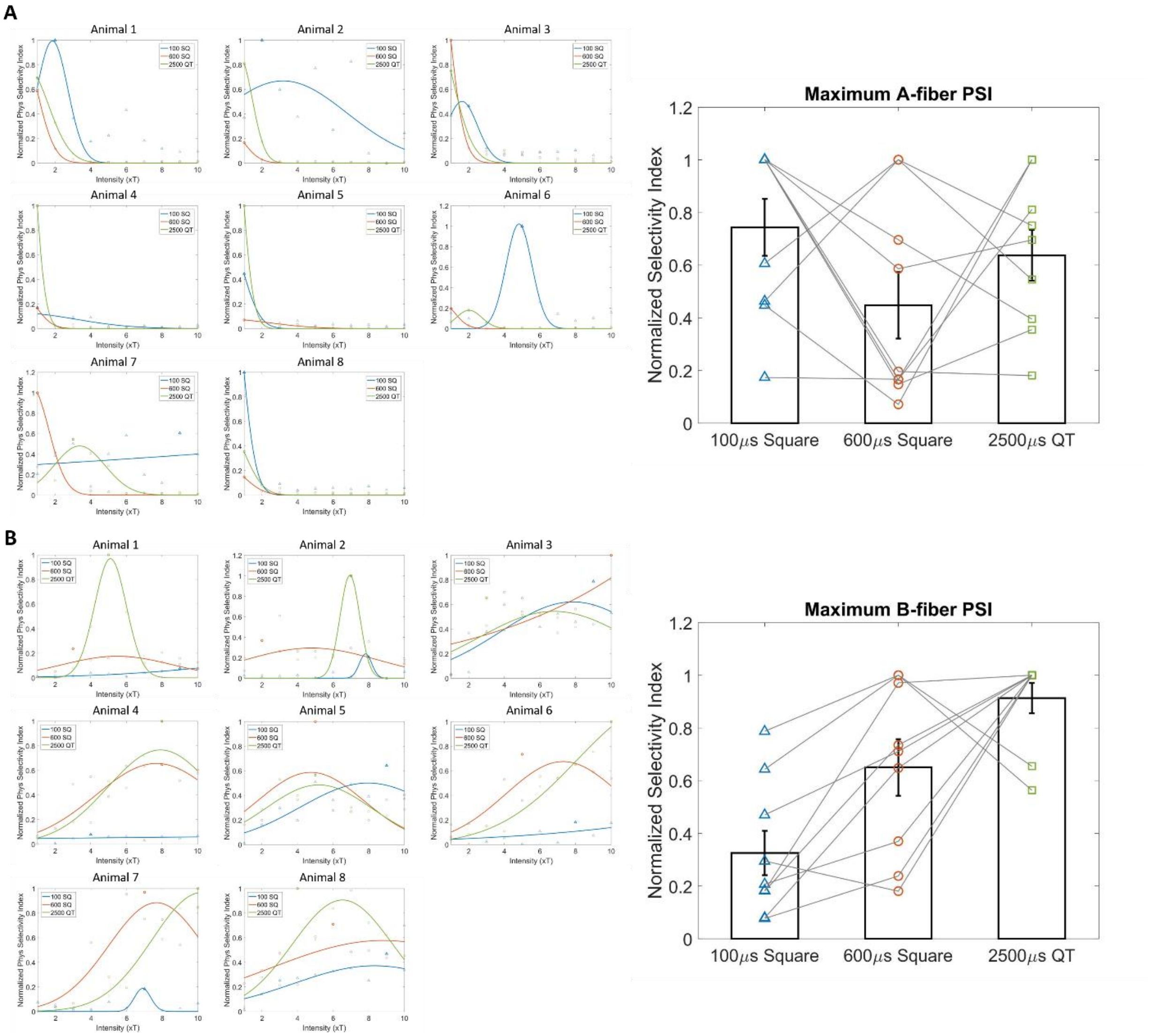


**Figure S10: The normalized A- and B-fiber PSIs in the rat model for different stimulation waveforms, including short square pulse (100µs), long square pulse (600µs), and proposed quasi-trapezoidal (100µs plateau+2500µs falling time). All the color/marker code are same as Figure 2.** (A) Left: The normalized A-fiber PSI for 8 animals. For each type of stimulus, the results were fitted by 1-d Gaussian function. Right: Comparison of maximum normalized A-fiber PSI for all stimulation parameters across animals (One-way ANOVA, $p=0.2$) (N=8). The dots with same color/marker code as (A) represent the maximum value of normalized A-fiber PSI for three types of stimuli respectively, and are linked as pair for each animal. The mean and standard error of are shown using bar charts with error bars (B) Same as (A) but for B-fiber (One-way ANOVA, $p<0.05$) (N=8).

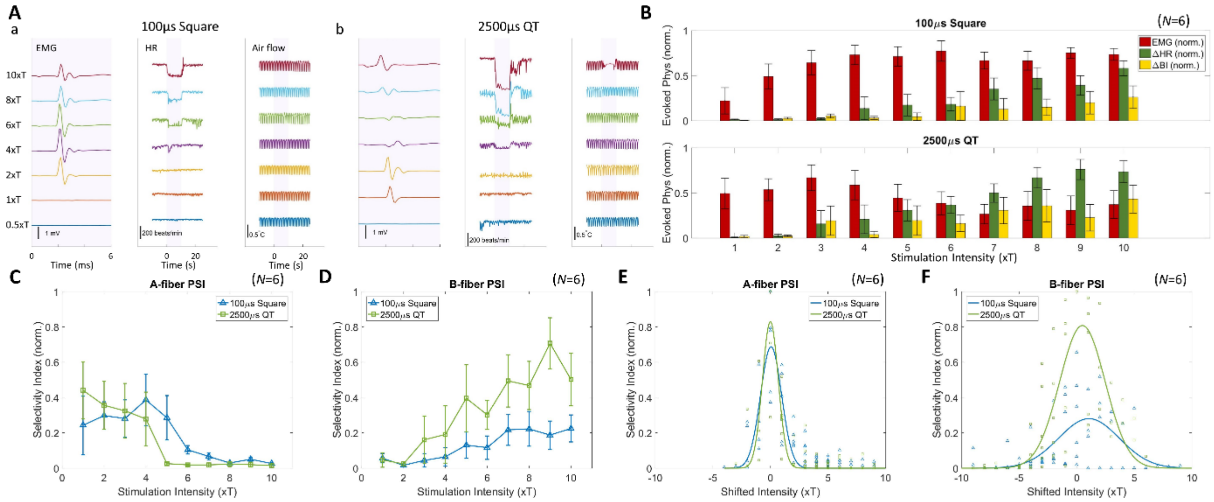


**Figure S11: VNS-elicited changes in physiological parameters, magnitude of physiological effects and resulting physiological selective indices (PSIs) in response to A- and B-fiber selective stimulus waveforms, in the mouse model.**

(A) Representative laryngeal EMG (left for each panel), heart rate (HR, middle of each panel), and breathing responses (air flow, right of each panel) elicited by stimulus trains of 2 different waveforms, short pulse (SP: panel a) and quasi-trapezoidal (QT: panel b), for a range of stimulus intensities (0.5 to 10×PT(EMG), close to T).
(B) Mean (±SE, N=6 animals) normalized EMG (red bars), HR (ΔHR, green bars) and breathing interval responses (ΔBI, yellow bars) elicited by the 2 stimulus waveforms, for different stimulus intensities. (ANCOVA, $p<0.05$ for waveform, across all physiological responses except for BI, $p<0.05$ for intensity, across all physiological responses except for EMG (0.73)).

(C) Mean (±SE, N=6 animals) normalized A-fiber PSI values associated with each of the stimulus waveforms (SP: blue, LP: red, QT: green trace), as a function of stimulus intensity. (ANCOVA, $p=0.55$ for waveform and $p<0.05$ intensity, $p>0.05$ for interaction).

(D) Same as (C) but for B-fibers. (ANCOVA, $p<0.05$ for waveform and intensity, $p<0.05$ for interaction). (E) Normalized A-fiber PSI values for each of the 2 waveforms plotted against stimulus intensity, re-aligned to the value associated with the maximum PSO for each animal and waveform, so the intensity shift of 0 corresponds the intensity producing the maximum PSI for each animal. Also shown are 1-D Gaussian function fits to the data for each of the 3 waveforms. Rmse for each fits: 0.134(SP), 0.128(QT). (ANCOVA, $p=0.86$ for waveform and $p<0.05$ for absolute intensity shift, p>0.05 for interaction). (F) Same as (E) but for B-fiber PSI values. Rmse for each fits: 0.140(SP), 0.224(QT). (ANCOVA, $p<0.05$ for waveform and absolute intensity shift, $p<0.05$ for interaction).

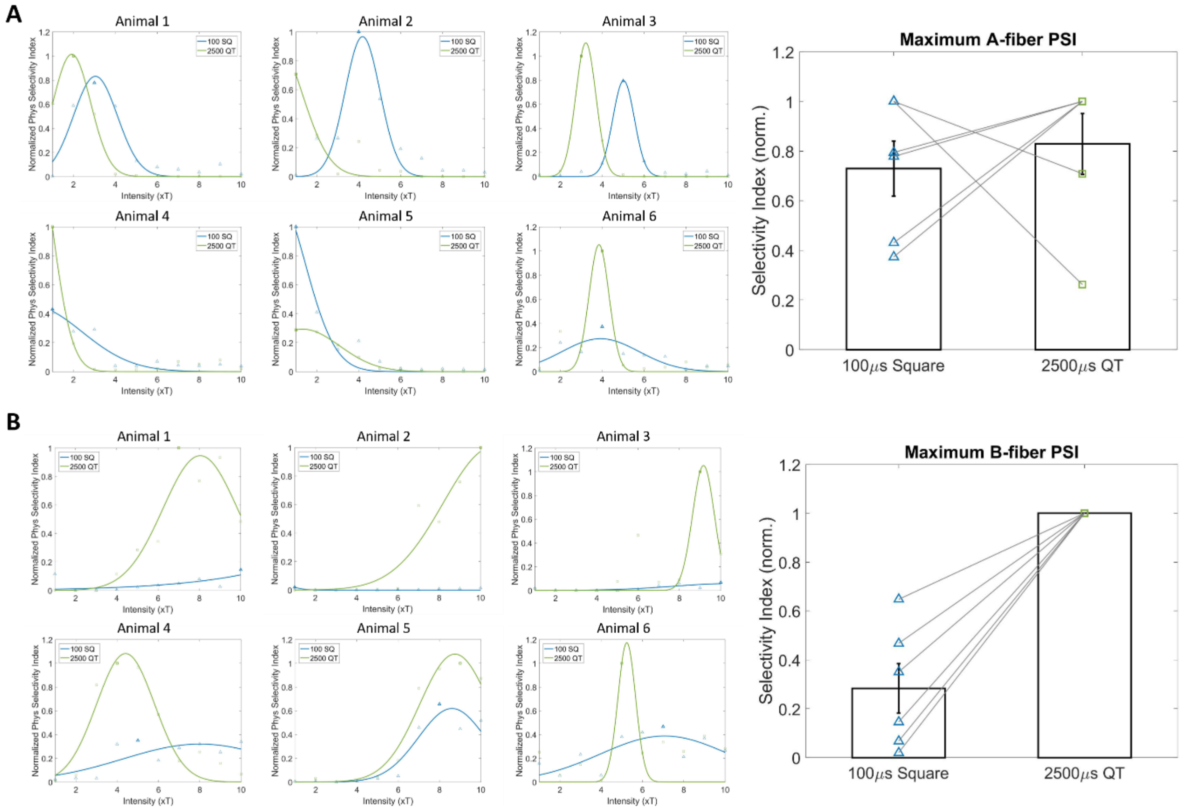


**Figure S12: The normalized A- and B-fiber PSIs in the mouse model for different stimulation waveforms, including short square pulse (100μs) and proposed quasi-trapezoidal (100μs plateau+2500μs falling time). All the color/marker code are same as Figure 3.** (A) Left: The normalized A-fiber PSI for 6 animals. For each type of stimulus, the results were fitted by 1-d Gaussian function. Right: Comparison of maximum normalized A-fiber PSI for all stimulation parameters across animals (One-way ANOVA, $p=0.54$) (N=6). The dots with same color/marker code as (A) represent the maximum value of normalized A-fiber PSI for three types of stimuli respectively, and are linked as pair for each animal. The mean and standard error of are shown using bar charts with error bars (B) Same as (A) but for B-fiber (One-way ANOVA, $p<0.05$).

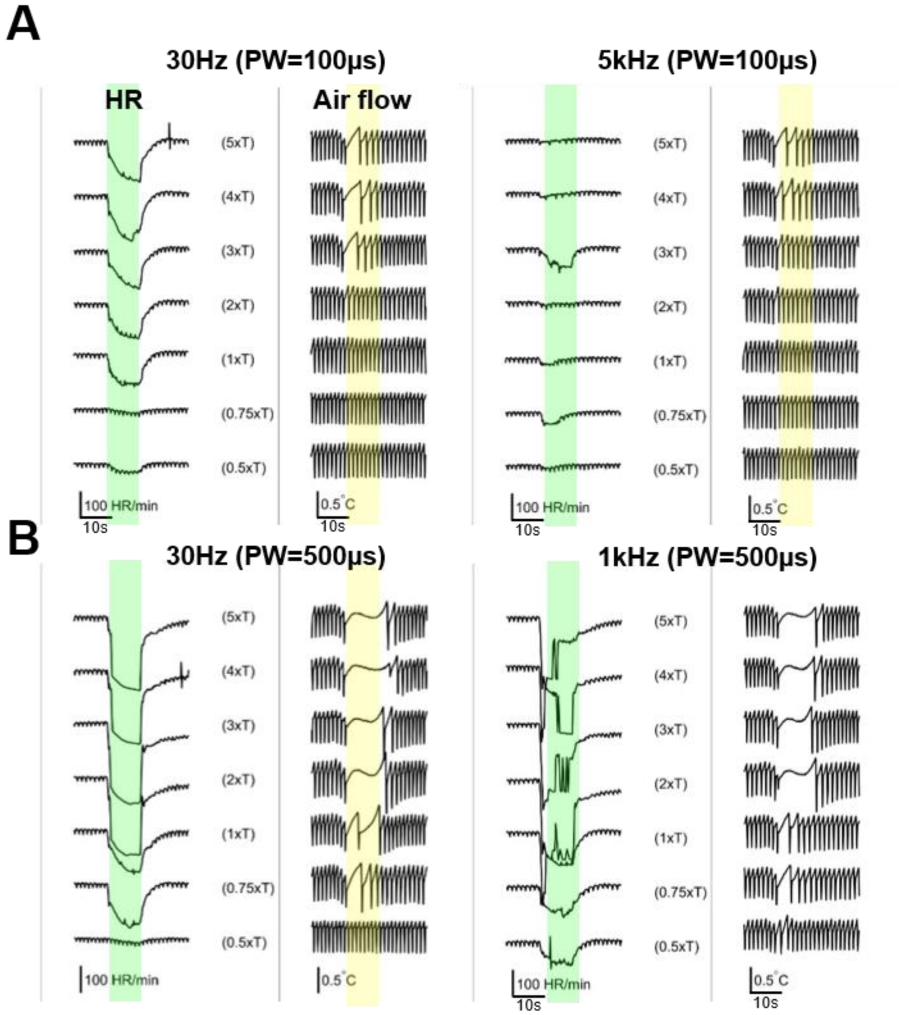


**Figure S13: VNS frequency modulates fiber selectivity**

(A) Representative heart rate (HR) and breathing responses (airflow) elicited by stimuli of 5kHz (100 µs PW), next to their 30 HZ, PW- and intensity-matched controls from the same animal.

(B) Same as (A) but for 1kHz (500 µs PW) and its corresponding 30 Hz controls.

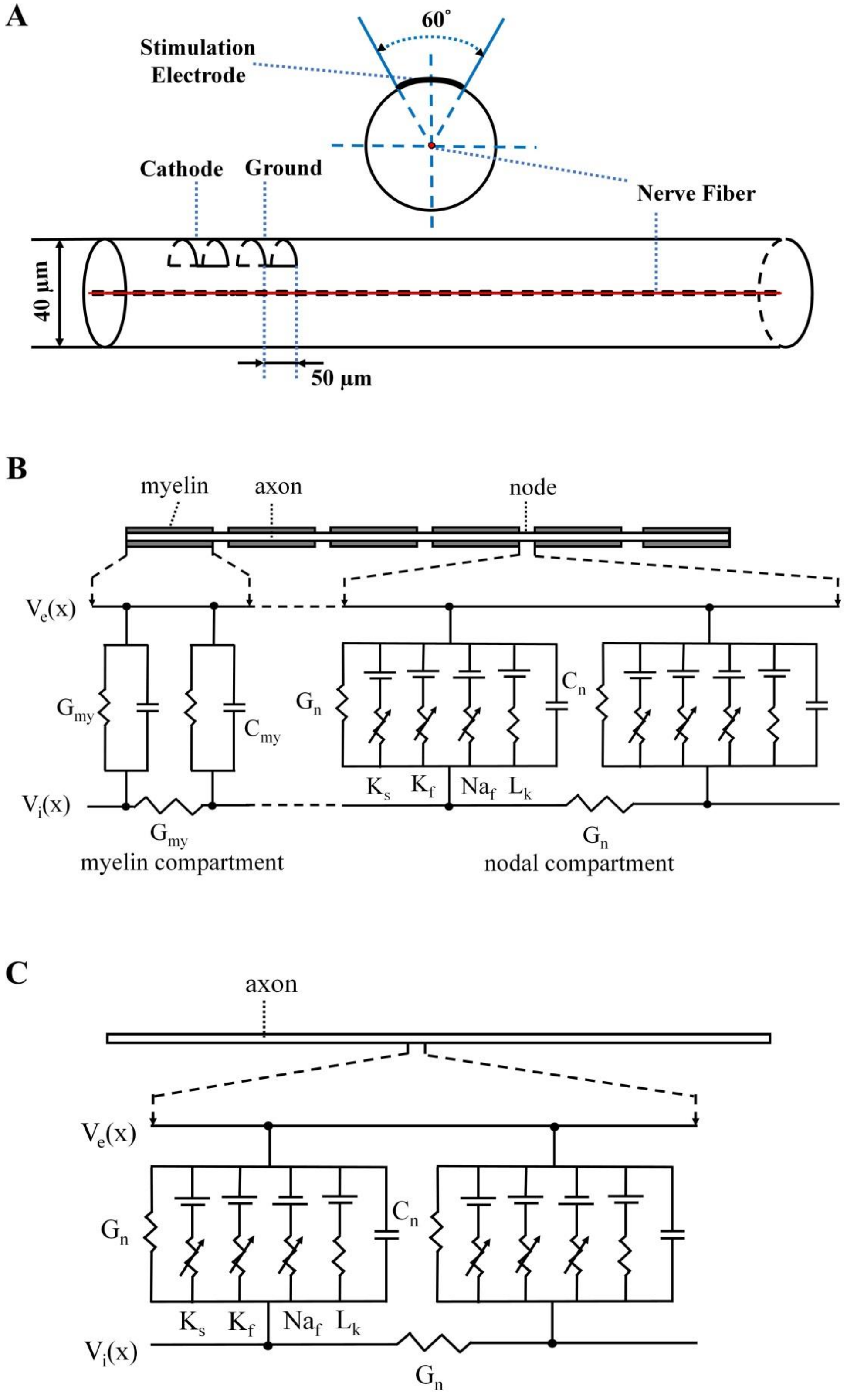
A
60°
Stimulation Electrode
Cathode
Ground
Nerve Fiber
40 μm
50 μm
B
myelin
axon
node
$V_e(x)$
$G_{my}$
$C_{my}$
$G_n$
$C_n$
$V_i(x)$
$K_s$ $K_f$ $Na_f$ $L_k$
$G_{my}$
myelin compartment
$G_n$
nodal compartment
C
axon
$V_e(x)$
$G_n$
$C_n$
$K_s$ $K_f$ $Na_f$ $L_k$
$V_i(x)$
$G_n$

**Figure S14. Schematic diagram of the fiber cable computational models and cuff stimulation electrode configuration.** (A) The nerve fiber (red) is defined as a 1D line at the center of the extracellular cylinder. Two 50-µm long cuff electrodes are located at the surface of the cylinder, with the electrode edges forming a 60° angle with the center. (B) Model structure of myelinated fibers. Myelin compartments were approximated by a linear conductance in parallel with a capacitor, whilst nodes were modelled by a parallel combination of three ion channel conductances ($Na^+$, fast $K^+$ and slow $K^+$) and a leakage conductance. (C) Model structure of unmyelinated fibers. $G_n$ and $G_{my}$ are the nodal and myelin conductivities respectively.

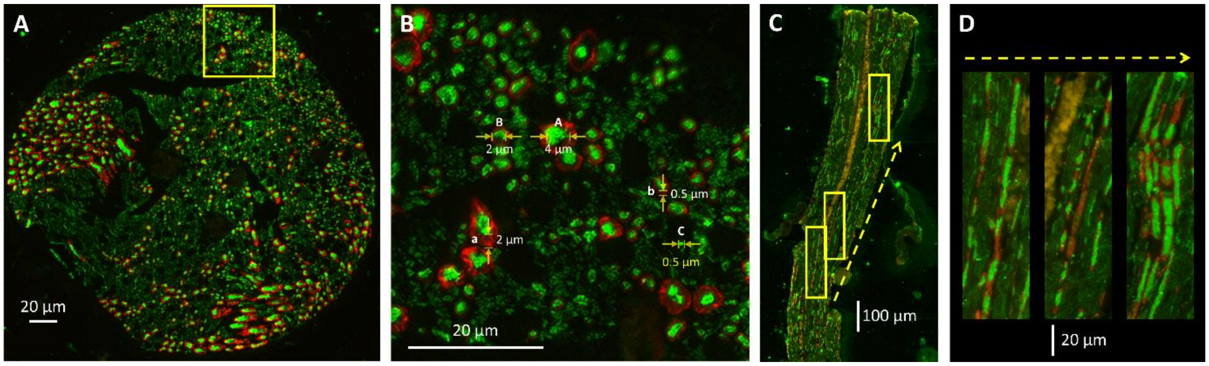


**Figure S15: Histology of rat cervical vagus nerve.** (A) Overview of cross sectional VN. Green and red fluorescence labelling represent the fiber (Neurofilament, AB8135, Abcam) and myelin structures (myelin basic protein, AB7349, Abcam) respectively. (B) Zoom-in cross sectional view and representative measurements of different types of fibers; A: Diameter of large myelinated fiber (A-type). a: Size of myelin of A-type fibers. B: Diameter of medium myelinated fibers (B-type). b: Size of myelin of B-type fibers. C: Diameter of small unmyelinated fibers (C-type). (C) Longitudinal sectional VN. (D) Zoom-in longitudinal sectional view and different lengths of myelin structure.

| Experiment | Recording type | Stimulation Configuration | Threshold type | Threshold range (µA) | # of animals |
|---|---|---|---|---|---|
| **Waveform Manipulation - Neural response** | Neural | Tripolar (rostral); Monophasic; Random pulse; | T (PW=100) | 10-40 | 5 |
| **Waveform Manipulation - Physiological response** | Neural+ Physiology | Tripolar (rostral) Monophasic; 10-s pulse train, frequency fixed at 30Hz | T (PW=100) | 12-50 | 8 |
| **Frequency Manipulation - High versus low frequency** | Physiology | Tripolar (caudal) Biphasic; 10-s pulse train, High frequency vs 30Hz | PT (PW=100) | 60-180 | 5 |
| **Frequency Manipulation - Multiple frequencies** | Physiology | Tripolar (caudal) Biphasic; 10-s pulse train, PW fixed at 40µs | PT (PW=40, 30Hz) | 150-300 | 5 |
| **Frequency Manipulation - KES + probing pulse** | Neural+ Physiology | Tripolar (caudal) Biphasic KES; Monophasic probing Pulse @ 1Hz; 10-s pulse train | PT (KES); NT (probing pulse: 100µs, 600µs) | KES: 200-250<br>100µs: 40-90<br>600µs: 30-50 | 7 |
| **c-Fos activity quantification** | Physiology | Tripolar (caudal) Biphasic KES (40 µs) vs 30Hz (100 µs) control; 30-mins intermediate stimuli, 10-s ON and 50-s OFF. | PT (KES) | KES: 200-250 | 12 |

**Table S1: Stimulation configuration, recording type, and thresholding method, threshold range, and number of animals for each rat experiment.**

| Experiment | Recording type | Stimulation Configuration | Threshold type | Threshold range (μA) | # of animals |
|---|---|---|---|---|---|
| **Waveform Manipulation - Physiological response** | Physiology | Tripolar (rostral) Monophasic; 10-s pulse train, frequency fixed at 30Hz | PT (EMG, close to T) (PW=100) | 2-13 | 6 |
| **Frequency Manipulation - KES** | Physiology | Tripolar (caudal) Biphasic; 10-s pulse train, PW fixed at 40μs | PT (PW=40, 30Hz) | 24-100 | 5 |

**Table S2: Stimulation configuration, recording type, and thresholding method, threshold range, and number of animals for each mouse experiment.**

| Model parameters | Value |
|---|---|
| Potassium equilibrium potential ($V_K$) | -84 mV |
| Unspecific ion equilibrium potential ($V_l$) | -84 mV |
| Nodal capacitance ($C_n$) | 2 μF/cm² |
| Myelin capacitance [a] ($C_{my}$) | 0.01 μF/cm² |
| Extracellular resistivity ($\rho_e$) | 75 Ω·cm |
| Nodal resistivity ($\rho_n$) | 75 Ω·cm |
| Maximum fast K+ conductance [b] ($g_{Kf}$) | 60.75 mS/ cm² |
| Maximum slow K+ conductance [b] ($g_{Ks}$) | 121.51 mS/ cm² |
| Nodal leakage conductance [b] ($g_l$) | 121.51 mS/ cm² |
| Sodium permeability [b] ($P_{Na}$) | 0.01426 cm/s |
| Extracellular sodium concentration [b] ($[\mathrm{Na}]_o$) | 154 mM |
| Intracellular sodium concentration [b] ($[\mathrm{Na}]_i$) | 35 mM |
| Faraday constant [b] ($F$) | 96485 C/mol |
| Gas constant [b] ($R$) | 8314.4 mJ/(K·mol) |
| Temperature [b] ($T$) | 310.15 K |
| **Myelinated fiber** | |
| **Axon diameter** | 2.6 μm |
| Myelin resistivity ($\rho_{my}$) | 65 Ω·cm |
| Myelin diameter | 5 μm |
| **Un-myelinated fiber** | |
| Axon diameter | 1.3 μm |

a. The myelin capacitance and resistance is estimated to match in vivo compound nerve action potential recordings in rats. b. Modified from Schwarz, Reid and Bostock (SRB) model.

**Table S3: Vagus nerve computational model parameters.**
